# Defining function of wild-type and patient specific *TP53* mutations in a zebrafish model of embryonal rhabdomyosarcoma

**DOI:** 10.1101/2021.04.21.440757

**Authors:** Jiangfei Chen, Kunal Baxi, Amanda E. Lipsitt, Nicole R. Hensch, Long Wang, Antoine Baudin, Daniel G. Robledo, Abhik Bandyopadhyay, Aaron Sugalski, Anil K. Challa, Andrea R. Gilbert, Gail E. Tomlinson, Peter Houghton, Eleanor Y. Chen, David S. Libich, Myron S. Ignatius

## Abstract

In embryonal rhabdomyosarcoma (ERMS) and generally in sarcomas, the role of wild-type and loss or gain of function *TP53* mutations remains largely undefined. Eliminating mutant or restoring wild-type p53 is challenging; nevertheless, understanding *TP53* effects on tumorigenesis remains central to realizing better treatment outcomes. In ERMS, >70% of patients retain wild-type *TP53*, yet *TP53* mutations when present in tumors are associated with poor prognosis. Employing a *kRAS^G12D^*-driven ERMS tumor model and newly generated tp53 null (tp53^-/-^) zebrafish, we define both wild-type and patient-specific *TP53* mutant effects on tumorigenesis. We demonstrate that *tp53* is a major suppressor of tumor initiation, where *tp53* loss expands tumors initiation from <35% to >97% of animals. Next, characterizing three patient-specific mutants finds that *TP53^C176F^* partially retains wild-type p53 apoptotic activity that can be exploited, while the *TP53^P153Δ^ and TP53^Y220C^* mutants define two structural mutations that predispose to head musculature ERMS.

## Introduction

*TP53* is the best-known tumor suppressor protein that is mutated or functionally disrupted in more than 50% of human tumors (Kastenhuber & Lowe, 2017; Muller & Vousden, 2014). Germline mutations in *TP53* are responsible for Li-Fraumeni Syndrome that predisposes to a wide but distinct spectrum of tumors that change with age (Malkin, 2011). Comprehensive analyses of *TP53* function *in vivo* and *in vitro* have revealed three different ways by which *TP53* can modulate tumorigenesis. These include effects caused by loss-of-function mutations, gain-of-function mutations, or dominant-negative mutations that disrupt the function of wild-type protein (Ko & Prives, 1996; Levine, Momand, & Finlay, 1991). Furthermore, mutant p53 function can be specific to cancer subtype (e.g., G>T transversions in lung cancer that do not correspond to the classic *TP53* hotspot mutations) (Olive et al., 2004; Petitjean, Achatz, Borresen-Dale, Hainaut, & Olivier, 2007). Recent studies comparing *TP53* mutation status with outcome in rhabdomyosarcoma patients show that *TP53* mutation status is correlated with increased risk and poor prognosis (Casey et al., 2020); however, the role for specific *TP53* mutations on sarcoma progression remains to be defined.

*TP53* mutations are present in >40% of adult carcinomas and thought to play a major role in tumorigenesis (Grobner et al., 2018). In contrast, with some exceptions, mutations in *TP53* are found in less than 6% of pediatric cancers (X. Chen et al., 2014; Chen et al., 2013; Grobner et al., 2018; Seki et al., 2015; Shern et al., 2014; Tirode et al., 2014). A major exception is Li-Fraumeni Syndrome, where germline *TP53* mutations predispose to a unique tumor spectrum that includes soft tissue sarcomas, such as rhabdomyosarcoma, and bone tumors (Guha & Malkin, 2017; Malkin, 2011). It is important to note that the *TP53* pathway is often suppressed in sarcomas. For example, in human embryonal rhabdomyosarcoma (ERMS), a common pediatric cancer of muscle, the *TP53* locus is mutated or deleted in 16% of tumors while transcriptional activity is altered in >30% of tumors, either through direct locus disruption or MDM2 amplification (Chen et al., 2013; Seki et al., 2015; Shern et al., 2014; Taylor et al., 2000). *TP53* mutations can also be acquired in ERMS at relapse, suggesting a role in tumor progression and/or resistance to therapy (Chen et al., 2013).

Mouse models have led the way in understanding *Tp53* function *in vivo*. Several murine genetic models were developed to assess the effects of both loss- and gain-of-function *Trp53* mutations (Attardi & Donehower, 2005; Garcia & Attardi, 2014) with mutant and null alleles both spontaneously developing cancer in multiple tissues (Lozano, 2010). Of note, the tumor spectrum in mice can vary depending on the mutant allele and genetic background; however, most *in vivo* studies have focused on a subset of hotspot mutations that are compared to the null or heterozygous wild type background and that are seen only in a subset of sarcoma patients (Attardi & Donehower, 2005; Garcia & Attardi, 2014; Guha & Malkin, 2017)). The vast majority of mutations observed in patients have not been interrogated in animal models, but rather function is inferred from these commonly studied hotspot mutants. *TP53* can have myriad effects on cancer biology and differences due to tumor type, mutation specific effects, or genetic background present significant challenges, with significant impact on therapeutic approaches in the clinic.

To define *tp53* biology *in vivo* using zebrafish, we recently generated a complete loss-of-function *tp53* deletion allele in syngeneic CG1-strain zebrafish using TALEN endonucleases (Ignatius et al., 2018). *tp53^del/del^* (*tp53^-/-^*) zebrafish spontaneously developed a spectrum of tumors that includes malignant peripheral nerve-sheath tumors (MPNSTs), angiosarcomas, germ cell tumors, and an aggressive natural killer cell-like leukemia (Ignatius et al., 2018). The role for *tp53* in self-renewal and metastasis of *kRAS^G12D^*-induced ERMS tumors was assessed using cell transplantation assays and revealed that *tp53* loss does not change the overall frequency of ERMS self-renewing cancer stem cells compared to tumors expressing wild-type *tp53* (Ignatius et al., 2018). In contrast, *tp53^-/-^* ERMS were more invasive and metastatic compared to *tp53* wild-type tumors, providing new insights into how *tp53* suppresses ERMS progression *in vivo*. By taking advantage of the fact that in zebrafish, similar to humans, *tp53* loss-of-function is not required for tumor initiation, we employ *tp53^-/-^* zebrafish in the present study to further assess the role for *tp53* on *kRAS^G12D^*-driven ERMS. We find that wild-type *tp53* affects ERMS proliferation with a smaller effect on apoptosis. Next, expressing wild-type human *TP53*, we find that *TP53* is functional in zebrafish and potently suppresses ERMS initiation *in vivo*. We also define the pathogenicity of two *TP53* mutations, one found in a patient with ERMS and the other a variant of unknown significance found in a teenager with an aggressive osteosarcoma at our clinic. We find that these two mutations have very different effects on tumor initiation, location, proliferation, and apoptosis in our zebrafish model. Taken together, these analyses highlight the zebrafish ERMS model as a powerful and high-throughput system that can be used to characterize the spectrum of common and rare *TP53* mutations in sarcoma patients in the clinic.

## Results

### *tp53* is a potent suppressor of ERMS initiation, growth and invasion

Whole genome/exome/transcriptome sequencing analysis revealed that a majority of primary ERMS tumors are wild-type for *TP53*. However, *TP53* loss, mutation, or MDM2 amplification accounts for pathway disruption in approximately 30% of ERMS (Chen et al., 2013; Seki et al., 2015; Shern et al., 2014). Furthermore, *TP53* mutation is associated with poor prognosis (Casey et al., 2020). In the zebrafish ERMS model, expressing the human *kRAS^G12D^* oncogene in muscle progenitor cells is sufficient to generate ERMS with morphological and molecular characteristics that recapitulate the human disease (Langenau et al., 2007). Although robust, in a *tp53* wild-type genetic background, the upper limit of tumor formation observed is 40% by 50 days post injection/fertilization (Langenau et al., 2007). To test whether *tp53* suppresses ERMS initiation *in vivo*, we generated tumors in *tp53* wild-type (*tp53^+/+^*) and *tp53^-/-^* mutant zebrafish and compared aspects of tumorigenesis, including ERMS onset and growth. Co-injection of *kRAS^G12D^* and *DsRed* under the *rag2* promoter resulted in approximately 34% of *tp53^+/+^* animals (n= 49/143 animals) with tumors by 60 days post injection. In stark contrast, >97% of *tp53^-/-^* animals displayed tumors (n=139/142 animals) by 60 days. *kRAS^G12D^*-induced *tp53^-/-^* ERMS displayed a very rapid onset, with the majority of tumors arising within the first 20 days of life (Figure 1A; *p*<0.0001). We assessed tumor size and number of tumors initiated per zebrafish and found that *tp53^-/-^* mutant ERMS displayed a significant increase in size (Figure 1B, D; *p* < 0.0001) and in the number of tumors per animal (Figure 1B, C; *p*=0.016), compared to *tp53^+/+^* animals. Assessment of histology showed no major differences between *tp53^+/+^* and *tp53^-/-^* ERMS, and tumors formed in the head, trunk, or tail similarly in both backgrounds (Figure 1E, F, Figure S1B). We assessed relative expression of *kRAS^G12D^* and its downstream effector *dusp4* in tumors from *tp53^-/-^* and *tp53^+/+^* zebrafish and confirmed similar expression levels of *kRAS^G12D^* and *dusp4* using quantitative qPCR (Figure S1A; n=3 tumors each, *p* = 0.8966 Unpaired t test with Welch’s correction). Taken together, our analysis suggests that *tp53* is a major suppressor of tumor initiation in RAS-driven zebrafish ERMS. We next assessed the role for *tp53* loss-of-function on ERMS tumor growth by following tumor-burdened animals over time. Similarly aged wild-type and *tp53^-/-^* tumors were imaged once a week over the course of 3 weeks. While wild-type ERMS grew large tumors that remained localized, tumors in *tp53^-/-^* ERMS grew larger in size and spread very rapidly past neighboring somites (Figure S1C-E; *p*=0.054 in week 1; *p*<0.0001 in week 2 and 3; Student’s t test). We also observed metastases at locations away from the primary tumors in *tp53^-/-^* zebrafish (Figure S1C, *tp53^-/-^* fish 3). The invasive metastatic behavior of primary *tp53^-/-^* tumors is consistent with our earlier analysis that found transplanted secondary and tertiary *tp53^-/-^* tumors are highly metastatic using an *in vivo* metastasis transplantation assay (Ignatius et al., 2018).

**Figure 1:**
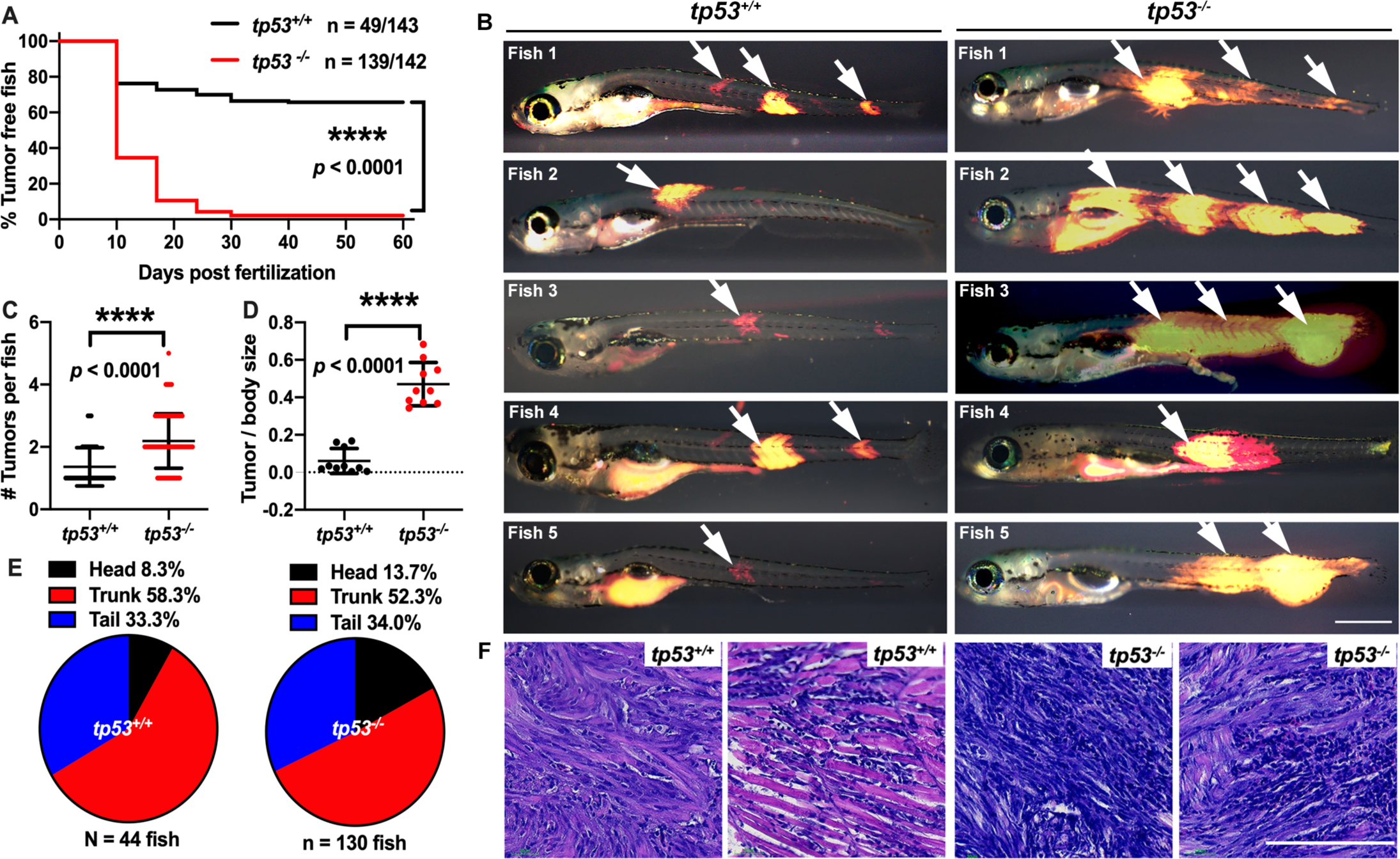
*tp53* suppresses ERMS tumor initiation. (A) Kaplan-Meier plot showing ERMS tumor initiation in *tp53^-/-^* and *tp53^+/+^* fish. (B) Representative images of DsRed-positive zebrafish ERMS. Arrows show tumor location for each fish. Scale bar = 1 mm. (C) Tumor numbers per zebrafish in *tp53^-/-^* and *tp53^+/+^* fish. n = 44 (*tp53^+/+^*), n = 130 (*tp53^-/-^*). (D) Ratio of tumor area to total body area in in *tp53^-/-^* and *tp53^+/+^* fish. n = 10. (E) Pie chart showing percentage of tumors found in varying regions of *tp53^-/-^* and *tp53^+/+^* fish, showing no significant differences in tumor localization. Head – *p* = 0.25848, trunk – *p* = 0.39532, tail – *p* = 0.92034 (Two-tailed Two Proportions Z test). (F) Representative H&E staining of zebrafish ERMS tumors. Scale bar = 100 µm.

### *tp53* suppresses proliferation and to a lesser extent apoptosis in ERMS tumors

Given that *tp53^-/-^* tumors are larger than equivalent stage wild-type tumors, we performed EdU and phospho-histone H3 staining to assess proliferation and Annexin V staining to assess apoptosis in primary ERMS. Treatment of wild-type and *tp53^-/-^* tumors with a 6-hour pulse of EdU revealed that *tp53^-/-^* tumors are significantly more proliferative (Figure 2A; *p* = 0.0002 Student’s T-test). Similarly*, tp53^-/-^* ERMS displayed significantly more phospho-histone H3-expressing cells when compared to *tp53* wild-type ERMS (Figure 2B; *p* < 0.0001 Student’s T-test). Analysis of Annexin V staining for apoptotic tumor cells showed no significant difference in the rate of late apoptosis (Q2) (Figure 2C-E; *p* = 0.592 Student’s t test), but rather a small yet significant decrease in early apoptosis (Q4) in *tp53^-/-^* mutant ERMS compared to wild-type (Figure 2C-E; *p* = 0.0073 Student’s T test). Taken together, our data suggests *tp53* is a potent suppressor of tumor cell proliferation, with a moderate effect on apoptosis during ERMS tumorigenesis.

**Figure 2:**
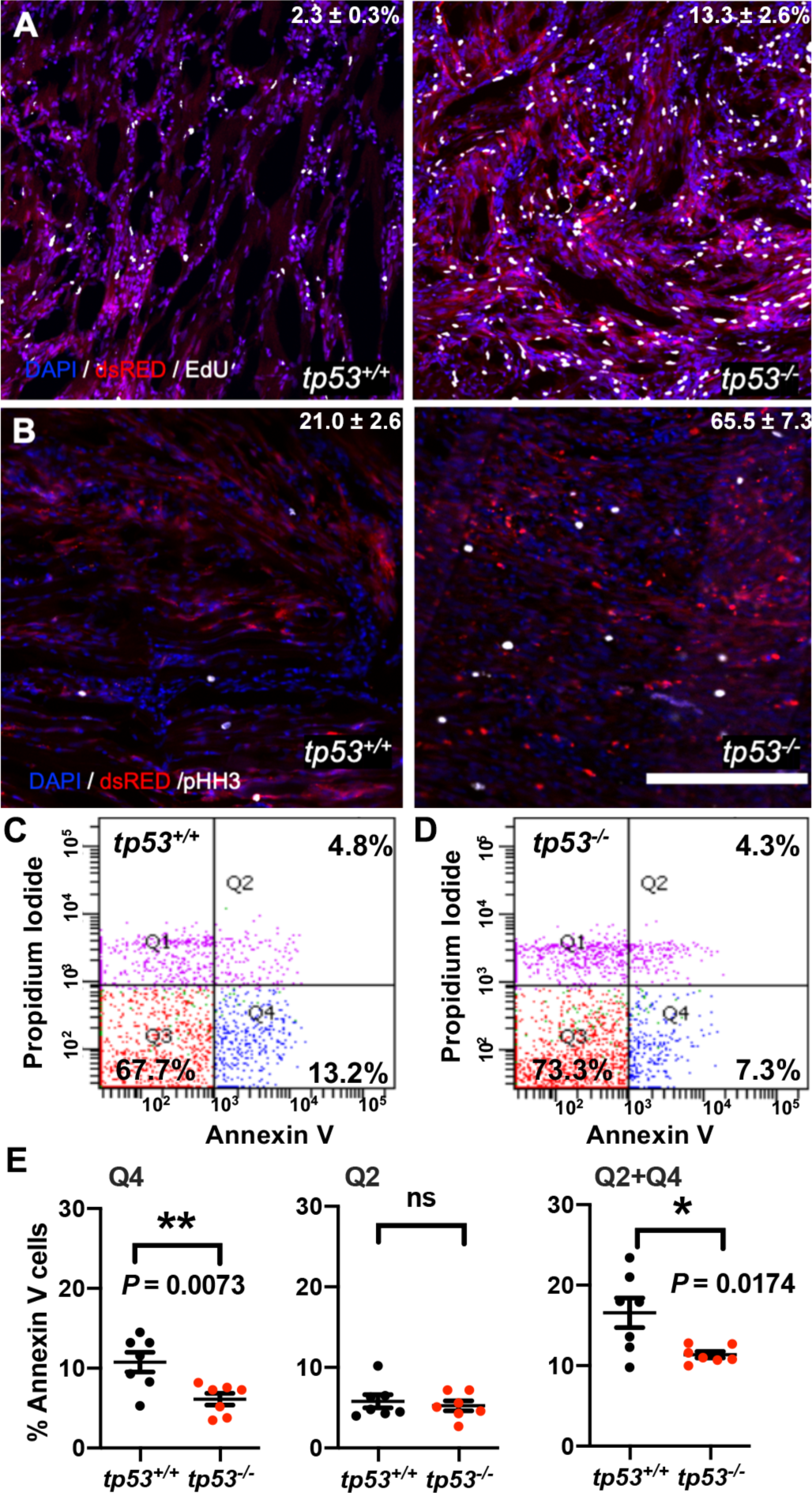
*tp53* is a potent suppressor of proliferation and to a lesser extent of apoptosis. (A) Representative confocal microscopy images of EdU staining on ERMS tumor sections n= 13∼16. (B) Representative confocal microscopy images of phospho-histone H3 staining on ERMS tumor sections. (Scale bar = 100 µm) n = 6. (C, D) Representative flow cytometry analysis of Annexin V staining of *tp53^+/+^*and *tp53^-/-^* ERMS tumors, respectively. (E) Quantification of flow cytometry analysis of Annexin V staining. Q1 = Pre-necrotic cells, Q2 = late apoptosis + necrotic cells, Q3 = living cells, Q4 = early apoptotic cells. n = 7. ns = not significant, *p* = 0.5926, Unpaired t test.

### Reintroducing human *TP53* in *tp53^-/-^* zebrafish blocks tumor initiation, growth, proliferation and increases apoptosis

We next assessed the consequence of introducing wild-type human *TP53* (*TP53^WT^*) in ERMS tumors in *tp53^-/-^* animals. Co-expression of *kRAS^G12D^* along with *TP53^WT^* significantly suppressed tumor initiation (Figure 3A; *p* < 0.0001, Student’s t test). ERMS in *TP53 ^WT^-*expressing animals grow more slowly and were unable to spread and/or metastasize (Figure 3B). The few tumors that did arise in *TP53 ^WT^*-expressing animals were smaller (Figure 3C; *p* < 0.0001 Student’s t test); however, there was no difference in the number or distribution of ERMS initiated per zebrafish between the two groups (Figure 3D, E; *p* = 0.065 for D, Student’s t test). Analysis of EdU staining of ERMS tumors determined that co-expressing *TP53^WT^* significantly inhibits proliferation (Figure 3F; *p* = 0.0007 Student’s t test). Also, we observed an overall increase in the number of apoptotic cells in *TP53^WT^*-expressing tumors as seen by Annexin V staining (Figure 3G; *p* < 0.0001 Student’s t test). We also assessed the effects of *TP53^WT^*on tumor initiation in the *tp53^+/+^* background and found no difference in the rate of tumor initiation for both groups (Figure S3A). Similarly, expression of wild-type zebrafish *tp53* by co-expression with *kRAS^G12D^* in *tp53^-/-^* zebrafish significantly suppressed tumor initiation (Figure S3B, C; *p* < 0.0001 Student’s t test). Furthermore, we found that this effect is dose-dependent, with co-expression of lower levels of *tp53* resulting in reduced suppression of tumor initiation in *tp53*^-/-^ zebrafish (Figure S3B; *p* < 0.0001 Student’s t test). Importantly, expression of human *TP53^WT^* or wild-type zebrafish *tp53* selectively in ERMS using the *rag2* promoter had no effect on the overall viability of zebrafish embryos or larvae.

**Figure 3:**
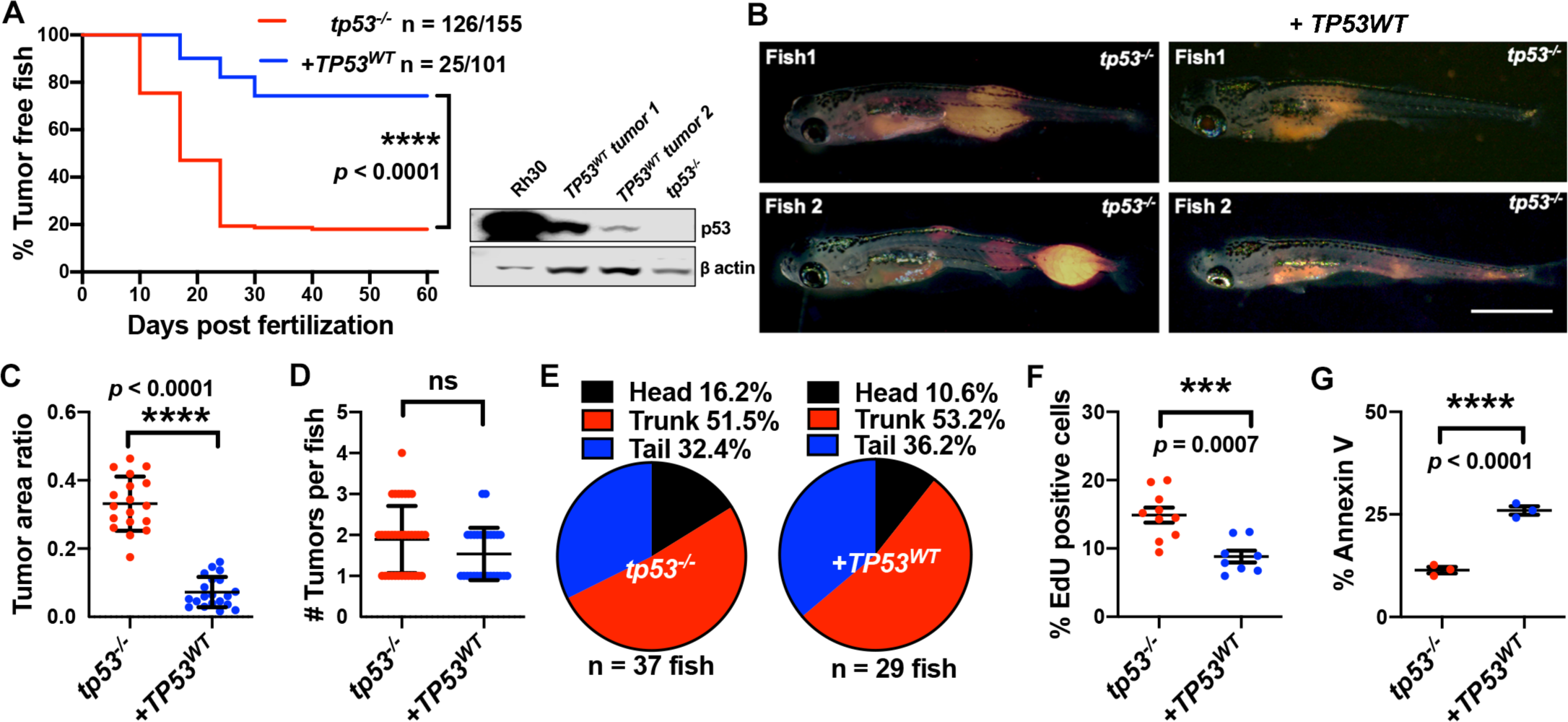
Human *TP53* blocks tumor initiation, growth, and proliferation and increases apoptosis in *tp53*^-/-^ zebrafish. (A) Kaplan-Meier plot showing ERMS tumor initiation in *tp53^-/-^* fish with or without p53^WT^ expression. Western Blot analysis was performed to assess p53^WT^ expression level in tumors. (B) Representative images of ERMS tumors in *tp53^-/-^* fish with or without human *TP53^WT^* expression. (C) Ratio of tumor area to total body area in *tp53^-/-^* fish with or without expression of *TP53^WT^*. n = 18. (D) Number of tumors per *tp53^-/-^* zebrafish with or without expression of *TP53^WT^*. ns = not significant. n = 36 (*tp53^-/-^*), n = 28 (*TP53^WT^*). (E) Pie chart showing site of tumor localization in *tp53^-/-^* fish with or without expression of *TP53^WT^* showing no statistical differences. Head – *p* = 0.20045, trunk – *p* = 0.42858, tail – *p* = 0.3336. Quantification of proliferation (F) and apoptosis (G) via EdU staining (n = 10) and Annexin V staining (n = 3), respectively, for tumors arising in *tp53^-/-^* fish with or without expression of *TP53^WT^*.

### Assigning pathogenicity to *TP53^P153Δ^* and *TP53^C176F^*, two undefined mutations in human sarcomas

The effects of specific *TP53* mutations on ERMS tumorigenesis are not defined. Analysis of *TP53* mutations in RMS patients found that a majority of mutations lie outside the well-studied hotspot locations and are mostly uncharacterized (Figure S4A). To determine if ERMS expressing specific human *TP53* point mutations differ from *TP53* deletion, we decided to study two uncharacterized mutations *in vitro* in SaOS2 osteosarcoma cells and *in vivo* by assessing mutant activity in ERMS generated in *tp53^-/-^* zebrafish. SaOS2 cells have the *TP53* gene deleted and provide a commonly used assay system to study effects of wild-type, hypomorphic, and gain-of-function effects in comparison to the *TP53^-/-^* background (Marcellus, Teodoro, Charbonneau, Shore, & Branton, 1996). The first mutation we selected is a *TP53^C^*^176F^ allele that is expressed by at least two patients with ERMS (Chen et al., 2013; Seki et al., 2015). In one of these patients, the second *TP53* allele was deleted in the tumor and the mutation is present in both the primary and relapsed tumor (Chen et al., 2013). The other mutation we selected to model is a rare *TP53^P153Δ^* (deletion of Proline 153, P153*Δ*) allele present in a patient in our clinic with an aggressive osteosarcoma. The patient developed an aggressive refractory osteosarcoma as a teenager, while her mother developed osteosarcoma in her early twenties; both succumbed to their disease. Based on the patient’s family history, tumor type, and *TP53* mutation, the diagnosis of Li-Fraumeni syndrome was rendered. However, since the mutation was rare, pathogenicity could not be assigned to this particular allele. Analysis of amino acid sequence conservation showed that residue C176 is conserved across humans, mouse, and zebrafish. On the other hand, P153 is present only in humans; however, it is located in a region that is highly conserved across all three species and the P153 is also conserved across other closely related mammalian species (Figure 4A, S4B). We confirmed the *TP53^P153Δ^* mutation by sequencing the DNA from the first passage patient-derived xenograft (PDX) generated from the patient at autopsy. Tumor cells obtained from the patient also harbored a missense mutation at c.476C>T (p.A159V) (Figure 4B).

**Figure 4:**
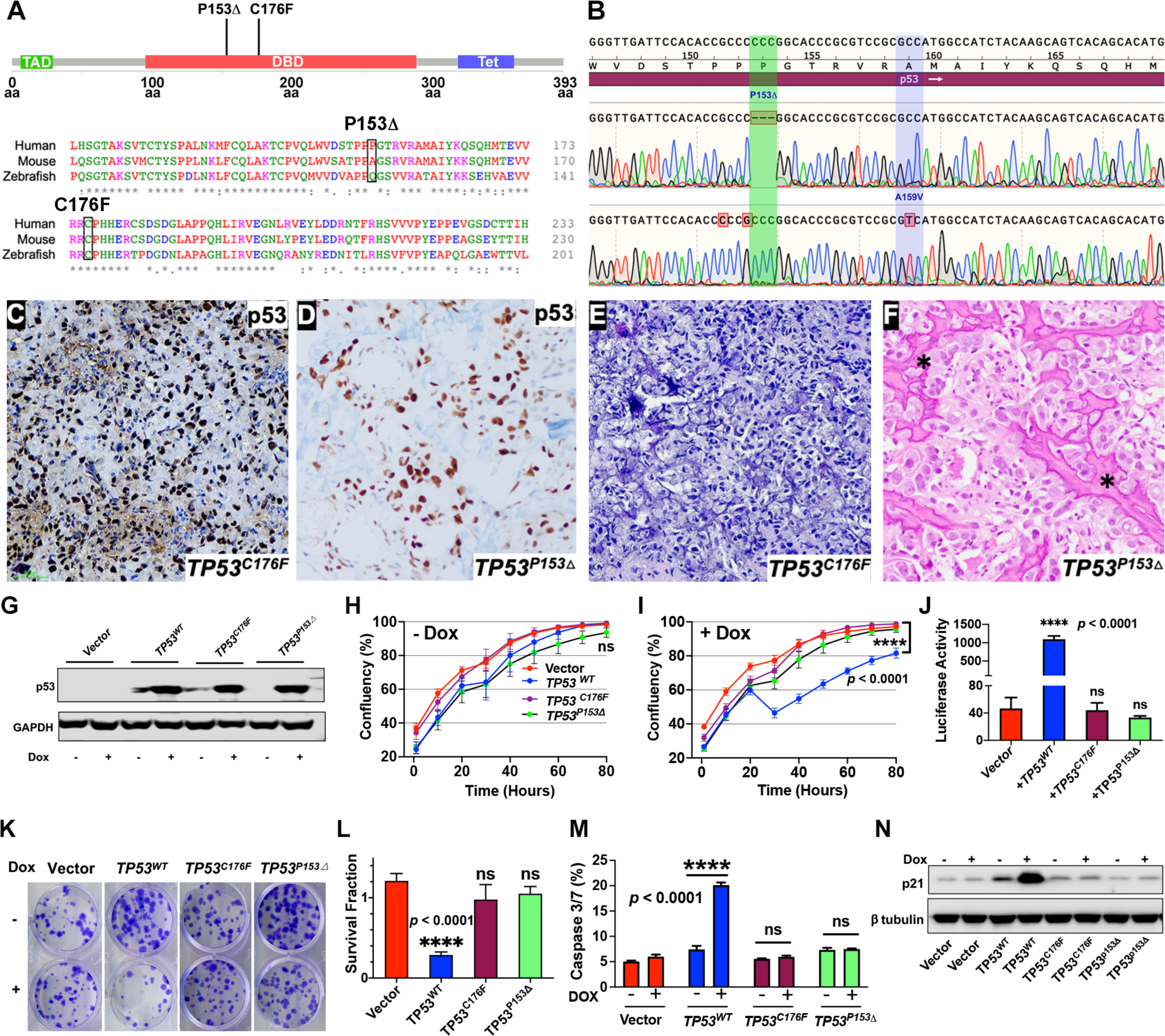
Assigning pathogenicity to two novel human *TP53* sarcoma mutations. (A) Lollipop plot showing the two novel, human *TP53* mutations P153*Δ* and C176F, as well as the amino acid sequence alignment for human, mouse, and zebrafish protein. (B) DNA sequencing data from osteosarcoma patient confirming the germline P153*Δ* mutation, as well as somatic A159V mutation. (C, D) p53 immunohistochemistry staining of p53 in ERMS PDX SJRHB00011 expressing p53^C176F^ and osteosarcoma expressing p53^P153Δ^. (E, F) Representative H&E staining of ERMS PDX expressing the C176F mutation and diagnostic biopsy of osteosarcoma tumor expressing osteosarcoma expressing p53^P153Δ^ showing neoplastic tumor cells with pleomorphic nuclei, irregular chromatin pattern, as well as irregular disorganized trabeculae of unmineralized malignant osteoid (stars). (G) Expression of p53^WT^, as well as p53^C176F^ and p53^P153Δ^ mutant protein, from a Dox-inducible pCW57.1 vector in SaOS2 osteosarcoma cells. (H, I) Growth curves (% confluence) of osteosarcoma SaOS2 cells harboring Dox-inducible pCW57.1 vector in the absence and presence of Dox, respectively. ns = not significant. (J) Luciferase assay to assess mutant p53 transcriptional activity. ns = not significant. (K) Colony forming assay in osteosarcoma SaOS2 cells harboring Dox-inducible pCW57.1 with either mutant or WT *TP53* in the absence or presence of Dox, respectively. (L) Quantification of change in colony formation in J. ns = not significant. (M) Quantification of apoptosis in osteosarcoma SaOS2 cells harboring Dox-inducible pCW57.1 with either mutant or WT *TP53* in the absence or presence of Dox. ns = not significant. (N) p21 expression in SaOS2 cells harboring Dox-inducible pCW57.1 with either WT or mutant p53 in the presence or absence of Dox.

We next assessed if the p53^C176F^ and p53^P153Δ^ proteins were expressed in the ERMS PDX SJRHB00011 that harbors the *TP53^C176F^* mutation and in the primary osteosarcoma that harbors the *TP53^P153Δ^* mutation using immunohistochemistry on tumor sections (Chen et al., 2013). Both the SJRHB00011 PDX tumor and the primary osteosarcoma tumor expressed p53 protein, as evidenced by strong positive nuclear staining in the majority of tumor cells (Figure 4C, D). Further H&E staining confirmed the rhabdomyosarcoma (heterogeneous population of ovoid to slightly spindled cells; Figure 4E) and osteosarcoma diagnosis (pleomorphic neoplastic tumor cells with irregular disorganized trabeculae of unmineralized malignant osteoid; Figure 4F).

To determine the effects of these mutations on tumorigenesis, we expressed either wild-type p53 (p53^WT^) or mutant p53^C176F^ or p53^P153Δ^ proteins in *TP53-*null SaOS2 osteosarcoma cells using a Doxycycline (Dox)-inducible vector (Figure 4G). Surprisingly, we find that only Dox-induced expression of wild-type p53 protein resulted in a reduction of cell growth (Figure 4H, I, *p* < 0.0001). We next assessed the ability of mutant and wild-type p53 proteins to activate transcription using a luciferase-based reporter containing 13 copies of the p53-DNA binding consensus sequence (el-Deiry et al., 1993), and again found that only wild-type p53 could drive significant transcriptional activity (Figure 4J, *p* < 0.0001). Next, we assessed the ability of mutant and wild-type p53 to suppress colony formation *in vitro*. Consistent with earlier results, expression of wild-type *TP53* reduced colony formation compared to empty vector controls; however, expression of *TP53^P153Δ^* and *TP53^C176F^* did not result in changes in colony formation compared to uninduced or SaOS2 control cells (Figure 4K, L, *p* < 0.0001, Dunnett’s multiple comparisons test). Using a Caspase-3/7 Glo apoptosis assay, we found that cells expressing wild-type p53 (+Dox) efficiently induced apoptosis. In contrast, uninduced control (-Dox) or cells expressing mutant p53 protein did not induce apoptosis (Figure 4M, *p* < 0.0001, two-way ANOVA with Sidak’s multiple comparisons test). Finally, compared to uninduced controls or cells expressing mutant p53 proteins, cells expressing wild-type p53 robustly induced p21 expression (Figure 4N). Thus, our *in vitro* results indicate that both *TP53* mutations (*TP53^P153Δ^* and *TP53^C176F^*) fail to retain wild-type function and do not show any gain-of-function activity in the SaOS2 *TP53^-/-^* background.

### *TP53^C176F^* is a hypomorphic allele while *TP53^P153Δ^* has gain-of-function effects in ERMS generated in *tp53^-/-^* zebrafish

We next assessed the two *TP53* mutations in ERMS generated in *tp53^-/-^* zebrafish. We co-expressed *TP53^C176F^* or *TP53^P153Δ^* along with *kRAS^G12D^* in *tp53^-/-^* embryos, as previously described. Expression of the mutant proteins in ERMS tumors was confirmed by Western blot analysis, using a human specific p53 antibody. For this assay, mutant p53 expression in Rh30 RMS cells was utilized as a positive control while zebrafish *tp53^-/-^* ERMS cells were used as a negative control (Figure 5A). Expression of *TP53^C176F^* with *kRAS^G12D^* in *tp53^-/-^* embryos resulted in a significant reduction in the initiation of tumors compared to expressing *kRAS^G12D^* alone in *tp53^-/-^* zebrafish (Figure 5B, *p* = 0.0005), while expression of *TP53^P153Δ^* along with *kRAS^G12D^* in the *tp53^-/-^* background resulted in similar rates of tumor initiation (Figure 5B, *p* = 0.774). We also assessed the effects of *TP53^C176F^* and *TP53^P153Δ^* on tumor initiation in the wild-type *tp53^+/+^* background and found no difference in the rate of tumor initiation for either of the mutant proteins (Figure 5C, *p* = 0.972).

**Figure 5:**
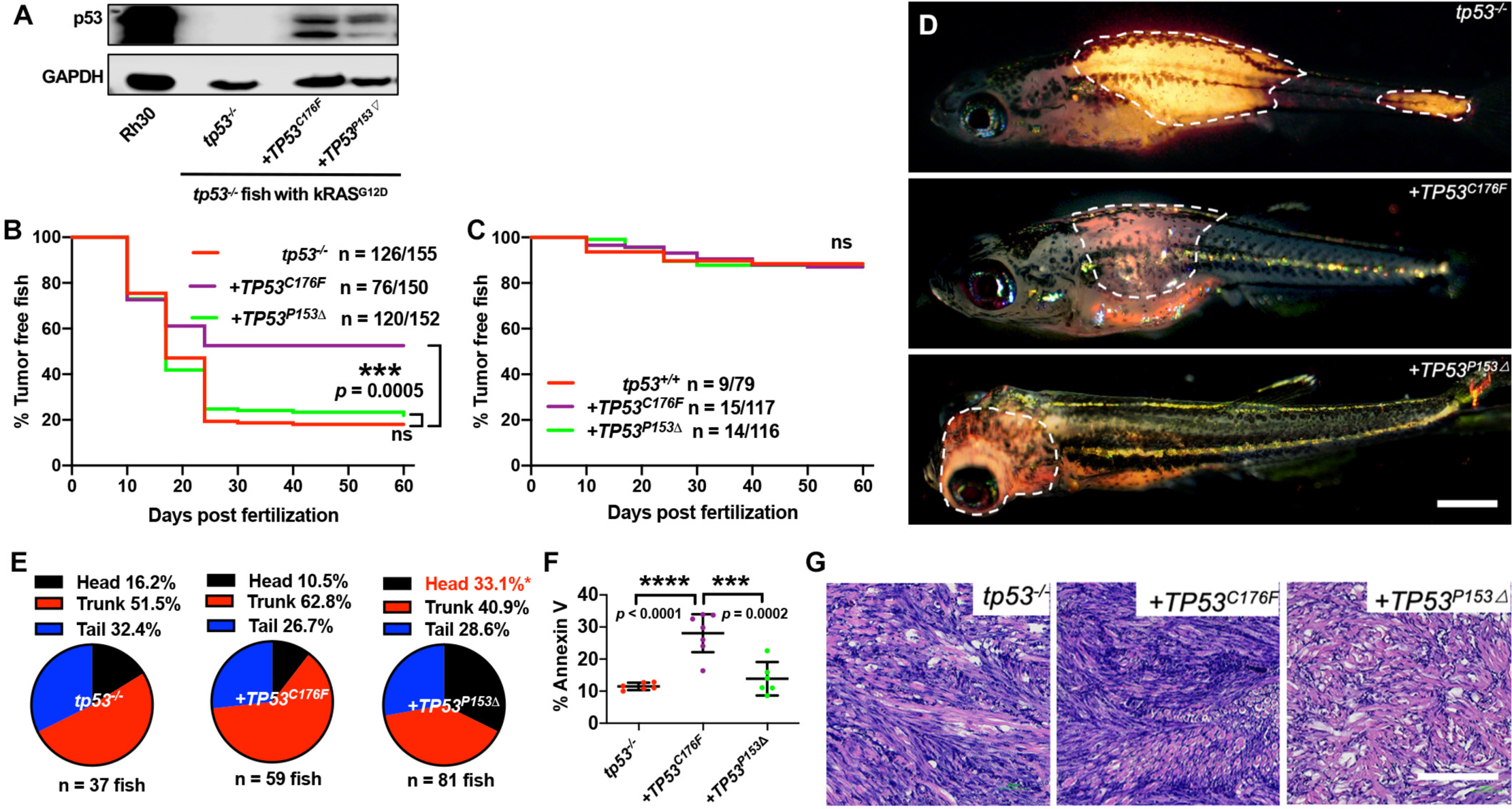
Assigning pathogenicity to two novel gain-of-function *TP53* mutations in *tp53^-/-^* zebrafish. (A) Protein expression of mutant p53 in zebrafish ERMS tumors, with RMS cell line, Rh30, as a control. (B) Kaplan-Meier plot showing tumor initiation in *tp53^-/-^* fish with or without expression of mutant *TP53*. (C) Kaplan-Meier plot showing tumor initiation in *tp53^+/+^* fish with or without expression of mutant *TP53.* ns = not significant (D) Representative images of tumor localization in *tp53^-/-^* fish with or without expression of mutant *TP53*. (E) Pie chart showing percentage of tumors found in varying regions of *tp53^-/-^* fish with or without expression of mutant *TP53.* Percentages in red indicate a significant difference to *tp53^-/-^* (*p* = 0.0096, Two-tailed Two Proportions Z test). (F) Quantification of Annexin V staining in tumors arising in *tp53^-/-^* fish with or without expression of mutant *TP53*. n = 6-7. (G) Representative H&E staining of tumors arising in *tp53^-/-^* fish with or without expression of mutant *TP53*.

Next, we assessed *TP53^C176F^* and *TP53^P153Δ^* -expressing tumors for tumor incidence and location. While the number of primary ERMS per fish in *tp53^-/-^* + *TP53^P153Δ^* were similar to *tp53*^-/-^ or *tp53^-/-^* + *TP53^C176F^*-expressing animals (Figure S5A, Versus *tp53^-/-^*, *p* = 0.065 (*TP53^C176F^*), *p* = 0.9299 (*TP53^P153Δ^*), one-way ANOVA with Tukey’s multiple comparisons test), rather unexpectedly, *tp53^-/-^* + *TP53^P153Δ^*-expressing zebrafish had greater than twice as many ERMS initiated in the head musculature (Figure 5D, E, *p =* 0.0096, Two Proportions Z test) indicating a gain-of-function effect with respect to site of tumor initiation.

Similarly, we assessed the effects of the *TP53* mutations on proliferation and apoptosis. The *tp53^-/-^* and *tp53^-/-^* + *TP53^C176F^*-expressing ERMS displayed increased levels of proliferation compared to *tp53^-/-^* + *TP53^P153Δ^*-expressing ERMS (Figure S5B, C, *p*=0.0062), while only *TP53^C176F^* tumors showed significantly higher rates of apoptosis, compared to the other two groups (Figure 5F, *p* = 0.0002). Tumor histology remained unchanged across all three groups (Figure 5G). Given that the *TP53^C176F^* mutation retains the ability to induce apoptosis, we next tested whether this activity could be augmented to inhibit tumor growth *in vivo*. It has been shown previously that ZMC1, a synthetic metallochaperone that transports zinc into cells as an ionophore, can restore p53 activity by stabilizing mutant p53 proteins such as p53^C176F^ (Blanden et al., 2015). To test this, we generated ERMS in the syngeneic CG1 strain zebrafish that were either *tp53^-/-^* or *tp53^-/-^ + TP53^C176F^*. Next, we expanded tumors from both groups in recipient wild-type CG1 animals, and treated tumors in recipient host animals for 2 weeks with either DMSO or ZMC1. Compared to DMSO-only control treatment, *tp53^-/-^ + TP53^C176F^* -expressing ERMS treated with ZMC1 showed a significant reduction in tumor growth over time (Figure S5D, G; *p*=0.0032 at week 3). We next assessed effects of ZMC1 on apoptosis via Annexin V staining and found that *tp53^-/-^ + TP53^C176F^* expressing tumors treated with ZMC1 displayed increased apoptosis after drug treatment (Figure S5F; *p* = 0.0008). Finally, we assessed p53^C176F^ expression and found that ZMC1 treatment increased protein levels, suggesting increased stability of p53^C176F^ protein (Figure S5E). In contrast, *tp53^-/-^* only ERMS treated with ZMC1 did not have any effect on the tumor growth (Figure S5H, K, *p* = 0.961) or apoptosis (Figure S5I-J, *p* = 0.583).

Altogether, we found that the *TP53^P153Δ^* mutation functions as a gain-of function allele with respect to the site of tumor initiation, while *TP53^C176F^* appears to retain some wild-type function, resulting in decreased ERMS initiation compared to tumor initiation in the *tp53^-/-^* background and the retention of the ability to trigger apoptosis *in vivo*. Additionally, the partial wild-type apoptotic activity of p53^C176F^ can be further enhanced by treatment with p53 reactivators, such as ZMC1.

### Expression of *TP53^Y220C^* predisposes to head ERMS in zebrafish

There is only one report of a patient with the *TP53^P153Δ^* mutation who had a predisposition to multiple cancers including breast cancer, soft tissue sarcoma and lung cancer (Michalarea et al., 2014). The location of the deleted proline suggests that the mutation causes a structural change in p53, therefore, we sought to understand the effect of this mutation on function by assessing protein structure and stability using *in silico* modeling. Homology models of p53^P153Δ^ generated by SWISS-MODEL (Waterhouse et al., 2018) indicate that deletion of P153 causes a partial narrowing of a small pocket on the surface of p53^WT^ (Figure 6A). P153 is the C-terminal residue of a tri-proline surfaced exposed loop that retains flexibility and mobility. Conversely, another known p53 mutation, the p53^Y220C^ mutation, causes an expansion and deepening of this pocket by forming a cleft bounded by L145, V147, T150-P153, P222, and P223 (Figure 6A). We searched the TCGA data base for other *TP53* mutants sequentially and structurally close to P153 and identified proline residues P151 and P152 are also mutated in cancers. Interestingly, we found mutations in P151 and P152 are present in a subset of tumors in the TCGA data base overlapping with the Y220C mutation (TCGA Research Network data; http://cancergenome.nih.gov). Previous molecular dynamics simulations of p53^Y220X^ (X = C, H, N or S) mutants found that the tri-proline loop (P151-153) is mobile and can precipitate a concerted collapse of the pocket, forming a frequently populated closed-state and contributing to the instability of p53 (Bauer et al., 2020). Since the binding of small molecules in the pocket increases the stability of p53^Y220X^ mutants, it is not inconceivable that mutations lining this pocket such as the tri-proline loop could be involved in fluctuations that decrease the structural stability of p53. Similarly, in our static models presented here, the loss of P153 causes this pocket to be slightly occluded and could contribute to further destabilization and effect p53 function. (Figure 6A). The *TP53^Y220C^* mutation is the ninth most frequent *TP53* missense mutation and is the most common “conformational” *TP53* mutation in cancer (Baud et al., 2018). Interestingly, the *TP53^Y220C^* allele is frequently associated with sarcomas and head and neck carcinomas and has been reported in patients with osteosarcoma and rhabdomyosarcoma (Castresana et al., 1995; Overholtzer et al., 2003). We therefore hypothesized that loss of P153 via an in-frame deletion might result in similar structural defects as the Y220C mutant by contributing to the overall structural instability of p53 (Wang & Fersht, 2017). Based on this hypothesis, we predicted that ERMS tumors expressing *TP53^Y220C^* may phenocopy *TP53^P153Δ^* with increased ERMS in the head region. To test this hypothesis, we generated ERMS that expressed *TP53^Y220C^* with *kRAS^G12D^* and *GFP* in *tp53^-/-^* embryos. Western blot analyses confirmed that mutant p53^Y220C^ protein was expressed (Figure 6B). Kaplan-Meier analyses indicated that while *TP53^Y220C^* expression inhibited ERMS initiation (Figure 6C, *p*<0.0001), decreased the number of primary ERMS per fish (Figure S6A, *p* <0.0001), *TP53^Y220C^* expression also led to a significant increase in head ERMS tumors compared to *tp53^-/-^* animals. This increase in head ERMS tumors was reminiscent of *tp53^-/-^* animals expressing *TP53^P153Δ^* (Figure 6D, F, H and Figure S6B; 31.1% of ERMS; n=150). We confirmed ERMS pathology by performing H&E staining on tumor sections (Figure 6E, G). As seen with *TP53^P153Δ^*, *TP53^Y220C^* expression led to a decrease in tumor cell proliferation (Figure 6J, *p*<0.0001); however, *TP53^Y220C^* expression also significantly increased apoptosis rates (Figure 6I, *p*=0.001), unlike that seen in *TP53^P153Δ^*-expressing ERMS.

**Figure 6:**
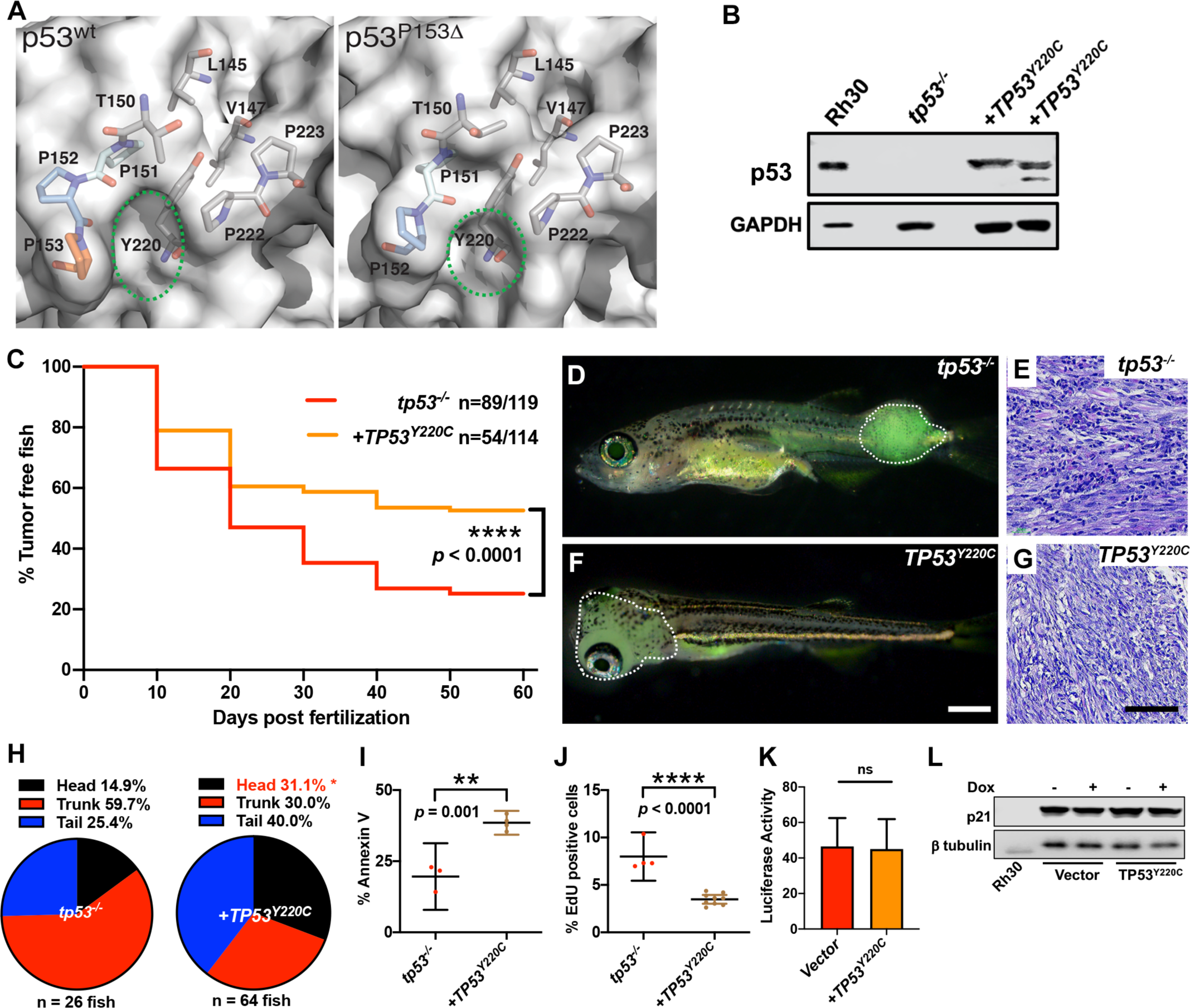
*TP53^Y220C^* predisposes to head ERMS in zebrafish. (A) Surface representation of p53^WT^ (PDB 2XWR) and p53^P153Δ^ (homology model) showing key residues lining a surface exposed pocket (sticks). The green ovals compare the size and shape of the pocket between the two structures. (B) p53 protein expression levels in *tp53^-/-^* fish tumors with or without *TP53*^Y220C^, with RMS cell line, Rh30, as a control. (C) Kaplan-Meier plot showing tumor initiation in *tp53^-/-^* fish, with or without *TP53*^Y220C^. (D) and (F) Representative images of *tp53^-/-^* fish with ERMS tumors, with or without *TP53*^Y220C^ (GFP positive). Dashed region outlines the tumor. (E) and (G) Representative H&E staining of tumors in *tp53^-/-^* fish, with or without *TP53*^Y220C^. (H) Pie chart showing localization of tumors expressed as a percentage found in varying regions of in *tp53^-/-^* fish with and without *TP53*^Y220C^. Percentage in red indicates a significant difference to *tp53^-/-^* (*p* = 0.01928, Two-tailed Two Proportions Z test). (I) Quantification of Annexin V staining in tumors of *tp53^-/-^* fish with or without expression of *TP53^Y220C^*. n = 3-4. (J) Quantification of EdU staining in tumors of *tp53^-/-^* fish with or without expression of *TP53^Y220C^*. n = 4-9. (K) Luciferase assay to assess mutant p53 transcriptional activity. ns = not significant. (L) p21 protein expression in SaOS2 cells harboring Dox-inducible pCW57.1 Vector only or expressing *TP53^Y220C^* in the presence or absence of Dox, with RMS cell line, Rh30, as a control.

We next assessed if *TP53^Y220C^* expression in SaSO2 cells displayed oncogenic activity. Similar to p53^C176F^ and p53^P153^*^Δ^*, p53^Y220C^ showed no discernible transcriptional activity compared to wild-type p53, as measured by the luciferase reporter assay and induction of p21 protein expression (Figure 6K, L, *p* =0.884). p53^Y220C^ also had no effect on SaOS2 growth or colony formation compared to wildtype p53 (Figure S6C-E, *p* = 0.283 (D, Ordinary one-way ANOVA with Tukey’s multiple comparisons test -Scr + Dox vs *TP53^Y220C^* + Dox), *p* = 0.1713 (E, Unpaired t test)). Finally, the p53^Y220C^ protein was unable to induce apoptosis when expressed using a Dox-inducible promoter and assessed by a caspase 3/7 apoptosis assay (Figure S6F, *p* = 0.5608, Welsh’s t test).

Altogether, our data reveal that both the *TP53^P153Δ^* and *TP53^Y220C^* mutations predispose to head ERMS tumors in zebrafish but differ in their effects on tumor initiation and apoptosis. However, similar to *TP53^C176F^* and *TP53^P153Δ^* mutants *in vitro* in SaSO2 cells, *TP53^Y220C^* does not show any effects on colony formation, transcriptional activity, proliferation, or apoptosis.

## Discussion

The *TP53* tumor-suppressor gene is mutated in >40% of human tumors, and patients with Li-Fraumeni syndrome with germ line mutations are predisposed to a spectrum of tumors that includes several potentially lethal childhood sarcomas, such as rhabdomyosarcoma and osteosarcoma (Grobner et al., 2018; Guha & Malkin, 2017). Here, we addressed three poorly understood aspects of *TP53* function in ERMS, a devastating pediatric malignancy of the muscle. First, we found that the *tp53* pathway is a major suppressor of tumor initiation in RAS-driven ERMS. Second, we found that human *TP53* can complement zebrafish *tp53* function. Third, we assigned function to two mutations whose effect in ERMS was previously unknown and defined *TP53^C176F^* and *TP53^P153Δ^* as a hypomorphic allele and pathogenic gain-of-function mutation, respectively. Lastly, we found that the structural mutations *TP53^P153Δ^ and TP53^Y220C^* predispose to head musculature ERMS.

Mutant RAS-driven ERMS tumors are associated with high-risk patients and are a challenge to treat in the clinic. RAS mutant ERMS models in mice are generated by introducing mutant RAS along with switching off *Trp53* and/or *p16* (Kashi, Hatley, & Galindo, 2015). In contrast, zebrafish ERMS is more similar to the human disease since RAS activation is sufficient to drive tumor formation. However, a maximum of 40% of wild-type *kRAS^G12D^*-expressing zebrafish initiate tumors by 50 days of life (Langenau et al., 2007). In both the zebrafish and human tumors with wild-type *tp53*, a subset of tumors have *MDM2* amplification and/or overexpression (Chen et al., 2013; Langenau et al., 2007; Seki et al., 2015; Shern et al., 2014). Additionally, other genes such as *TWIST1* have also been shown to promote tumor initiation in ERMS and sarcomas through suppression of p53 expression (Piccinin et al., 2012). Of note, in the zebrafish ERMS model, it was previously shown that incidence was increased to approximately 70% of animals in the *tp53^M214K/ M214K^* mutant background (Berghmans et al., 2005; Langenau et al., 2007), suggesting that *tp53* plays an important role in suppressing ERMS initiation; however other pathways may also function to initiate RAS-driven ERMS in the remaining 30% of animals that remain tumor free. However, it is important to note that the recently generated *tp53^-/-^* complete loss-of-function mutant zebrafish line (Ignatius et al., 2018) spontaneously form a broader tumor spectrum than the *tp53^M214K/ M214K^* allele, which includes MPNSTs, angiosarcomas, NK-cell leukemias, and germ cell tumors (Berghmans et al., 2005; Ignatius et al., 2018). Our study tests the role for complete loss of *tp53* in ERMS initiation by generating *kRAS^G12D^*-induced ERMS in the *tp53^-/-^* background. We found that in the *tp53^-/-^* complete loss-of-function background >97% of animals form tumors, indicating that *tp53* pathway suppression is required for ERMS full tumor penetrance. Moreover, when compared to control wild-type animals, *tp53^-/-^* animals initiate significantly more tumors per animal, display increased tumor cell proliferation, and exhibit a relatively less pronounced effect on tumor cell apoptosis. The effects on apoptosis can be enhanced through expression of either wild-type human *TP53* or zebrafish *tp53* under the control of the *rag2* promoter, specifically in ERMS tumor cells. Our study also demonstrates that in primary tumors, *tp53* is a potent suppressor of tumor cell growth, invasion, and metastasis, confirming earlier studies that used transplanted tumor cells into unlabeled syngeneic hosts (Ignatius et al., 2018). Our previous study also showed that loss of *tp53* had no effect on tumor cell self-renewal (Ignatius et al., 2018). Thus, while *TP53* is shown to suppress tumor progression by multiple mechanisms depending on tumor type, our results suggest that loss of *TP53* in human ERMS may increase aggressiveness through enhanced proliferation, invasion, and metastasis.

A second important finding is that human *TP53* can complement wild-type *tp53* function in zebrafish ERMS, which allows for the study of different human alleles directly *in vivo*. Moreover, studying the effects of different levels of *TP53* specifically in ERMS tumors does not drastically alter tumor phenotypes observed. The biggest difference observed was a slight increase in total Annexin-positive apoptotic cells; however, it is important to note that co-expression *in vivo* of mutant and wildtype p53 results is protein expression at comparable or at lower levels than present in human tumor cells that endogenously express mutant p53 protein. For example, our assays displayed lower p53 expression than that observed in Rh30 RMS cells, which express mutant p53^R273C^ (Gibson, Harwood, Tillman, & Houghton, 1998). Secondly, the co-expression approach we use has been applied to study multiple aspects of tumorigenesis in zebrafish including in ERMS, T-cell acute lymphoblastic leukemia (T-ALL), melanoma, liver cancer, and neuroblastoma (Blackburn et al., 2014; Langenau et al., 2007; Lobbardi et al., 2017; White, Rose, & Zon, 2013). In zebrafish ERMS, co-expression approaches are used to study the effects multiple pathways have on tumorigenesis, including Notch signaling, canonical and non-canonical Wnt signaling, Myf5 and MyoD on tumorigenesis; moreover many of the effects seen were validated using human cell culture or *in vivo* in murine xenograft assays (E. Y. Chen et al., 2014; Hayes et al., 2018; Ignatius et al., 2017; Tenente et al., 2017).

Having defined that wild-type human *TP53* indeed functions similarly to endogenous zebrafish *tp53*, we were able to assess three mutant forms of *TP53* observed in sarcomas and show they have very different effects on tumorigenesis. The *TP53^C176F^* mutation was identified in a patient with ERMS, with loss of the second *TP53* wild-type allele (Chen et al., 2013). A second *TP53^P153Δ^* mutation was found to be germline in a patient with osteosarcoma. Finally, the *TP53^Y220C^* mutation is commonly found in sarcomas, including RMS and osteosarcoma (Castresana et al., 1995; Overholtzer et al., 2003). Using our *in vivo* assays, we were able to identify that the *TP53^C176F^* allele functions as a hypomorph with respect to ERMS initiation that retains the ability to induce apoptosis. The trunk is a common site of tumor initiation for ERMS in zebrafish, and there was a slight predisposition for the *TP53^C176F^* allele to initiate tumors in the trunk. These findings are consistent with studies showing that the *TP53^C176F^* allele does retain some aspects of wild-type p53 function, can form tetramers with wild-type p53, and can differentially induce *TP53* target genes like *NOXA*, *P53R2, GADD45, BAX,* and *WAF1* (Hoffman-Luca et al., 2015; Kato et al., 2003). Surprisingly, the *TP53^C176F^* allele had no effect on colony formation when expressed in SaOS2 osteosarcoma cells, suggesting that this function was not maintained in this commonly used assay system. Our results are consistent with other reports showing that certain mutations that modulate stability of p53 often show a range of effects *in vitro* assays and altering the temperature in which the cells are grown can be used to study their function (Di Como & Prives, 1998; Pfister & Prives, 2017). Significantly, compounds like ZMC1 enhance p53-induced tumor cell apoptosis. ZMC1 is thought to stabilize mutant p53 protein through increasing Zn^2+^ ions in cells and decrease tumor growth (Blanden et al., 2020; Blanden et al., 2015; Yu, Vazquez, Levine, & Carpizo, 2012). We found that ZMC1 effectively stabilizes mutant p53^C176F^ and selectively increases ERMS apoptosis in *tp53*^-/-^ + *TP53^C176F^* but not *tp53^-/-^* tumors, revealing an effective *in vivo* drug efficacy platform to identify therapeutic strategies for p53 mutant ERMS in the future. Together, our data shows that in the context of ERMS, the *TP53^C176F^* allele is likely hypomorphic and stabilizing the resulting mutant protein can be used to increase apoptosis in the context of treatments.

In contrast to *TP53^C176F^*, *TP53^P153Δ^* is rare with one known patient reported in the literature (Michalarea et al., 2014). The patient was diagnosed with multiple tumors over several decades including bilateral breast cancer, malignant fibrous histocytoma, and an EGFR mutant lung adenocarcinoma. Several members of the patient’s family including her mother, maternal aunts, and maternal grandmother all died due to early onset cancers, meeting the criteria of a Li-Fraumeni diagnosis (Michalarea et al., 2014). The specific Proline 153 residue in human p53, while in a region that is highly conserved across species, is not conserved in mice or zebrafish, presenting a challenge for modeling *in vivo*. Nevertheless, the Proline 153 residue is conserved between human and closely related mammalian species. Due to its rarity in the literature and the lack of animal models, genetic testing done through Invitae, a medical genetic testing provider, could not assign pathogenicity to *TP53^P153Δ^*; however, the genetics strongly suggest that this is a Li-Fraumeni mutation based on both the patient and her mother being predisposed to osteosarcoma at young ages. Moreover, the experience in the clinic was that the osteosarcoma in the patient was extremely aggressive and refractory to multiple anti-cancer agents. Notably, our *in vivo* studies found that *TP53^P153Δ^* predisposes to head ERMS and that tumors are relatively resistant to apoptosis. Presently, no other gene or pathway has been identified in zebrafish that predisposes to head tumors, and head ERMS tumors are infrequent compared to tumors in the trunk and tail. Moreover, the *TP53^Y220C^* mutant in zebrafish also predisposed to head ERMS, revealing two *TP53* mutant proteins that have shared gain-of-function effects in ERMS, possibly suggesting that they may gain shared activity that predispose to head ERMS tumors.

Head and neck rhabdomyosarcomas form a significant subset of ERMS tumors, and currently the basis for predisposition to head and neck RMS tumors is not understood. In mice, expressing activated Smoothened protein in cells expressing ap2 results primarily in head and neck ERMS (Hatley et al., 2012). Smoothened is a key component of the Hedgehog signaling pathway that is commonly activated in ERMS (Drummond et al., 2018; Satheesha et al., 2016). However, cell of origin likely plays a major role, given that in an independent mouse model, activated Smoothened under the control of a more ubiquitously expressed promoter leads to ERMS tumors in other skeletal muscle populations (Mao et al., 2006). Our data suggests that differential features of *TP53* alleles like *TP53^P153Δ^* and *TP53^Y220C^* may also predispose patients to head ERMS, providing additional mechanistic insights into anatomical differences in sarcoma initiation. Further, human *TP53* mutations and loss of *tp53* in our zebrafish models clearly show that loss or mutant *TP53* is associated with enhanced tumorigenic properties, suggesting a reason some head and neck RMS tumors may be typically more aggressive in nature.

In summary, our study reveals the zebrafish ERMS model to be an effective tool to define multiple aspects of wild-type *TP53* tumor suppressor function and to delineate both partial loss-of-function and gain-of-function mutational effects *in vivo*. This is an especially useful tool for sarcomas where a majority of *TP53* mutations remain uncharacterized. Different *TP53* alleles identified in patient tumors have very different effects on tumorigenesis *in vivo* and can potentially respond differently to therapeutic compounds. Thus, the type of precision modeling demonstrated here promises to help further define patient-specific *TP53* biology and improve clinical strategies in the future.

## Materials and Methods

### Animals

Animal studies were approved by the UT Health San Antonio Institutional Animal Care and Use Committee (IACUC) under protocol #20150015AR (mice) and 20170101AR (zebrafish). Zebrafish strains used in this work include: AB wildtype, AB/*tp53^-/-^*, CG1 wild type, CG1/ *tp53^-/-^*, Tg casper; *myf5:GFP*. Zebrafish were housed in a facility on a continuous flow system (Aquarius) with temperature-regulated water (∼28.5°C) and a 14-hour light-10-hour dark cycle. Adult fish were fed twice daily with brine shrimp, supplemented with solid food (Gemma). Larval fish were kept off the continuous flow system until 15 days post-fertilization and supplemented with a paramecium/algae culture before transferred online.

### Generation of zebrafish rhabdomyosarcoma tumors

*rag2-kRASG12D* and *rag2-DsRed* plasmids were linearized with XhoI, followed by phenol-chloroform extraction and ethanol precipitation. The purified DNA was resuspended in nuclease-free water (AM9916, ThermoFisher) and injected into embryos at the one-cell stage of development (40 ng/μl of *rag2-kRASG12D* and 20 ng/μl of *rag2-DsRed*). For expression of zebrafish *tp53* or human *TP53* in ERMS tumors in zebrafish, XhoI linearized *rag2-kRASG12D* and *rag2-DsRed* plasmids were injected along with Xmn1 linearized *rag2-TP53* (human) or *rag2-tp53* (zebrafish) constructs. For expression of mutant human *TP53* experiments, tumors were generated by injecting XhoI linearized *rag2-kRASG12D* and *rag2-DsRed* or *rag2-GFP* along with Xmn1 linearized *rag2-TP53^C176F^*, or *TP53^P153Δ^*, or *TP53^Y220C^* into the one cell stage of zebrafish embryos <1 hour post-fertilization (Ignatius et al., 2012; Langenau et al., 2007). Animals were allocated into experimental groups based on which injection cocktail they received during microinjection. Animals were monitored for tumor onset beginning at 10 days post-fertilization by scoring for DsRed or GFP fluorescence under an Olympus MVX10 stereomicroscope with an X-Cite series 120Q fluorescence illuminator. Scoring for tumor initiation was conducted for 60 days. The tumor onset was visualized by Kaplan-Meier plot, and the significance was analyzed by Log-rank (Mantel-Cox) test for each two groups at a time. Sample size was determined taking into account the observed tumor initiation rates from previous studies using the zebrafish ERMS model (Langenau et al., 2008; Tenente et al., 2017).

### Tumor size (ratio) measurements

Tumor size was measured using FIJI image analysis software (https://fiji.sc/) by comparing area occupied by the tumor region to the total body area of the fish. Specifically, all tumor-burdened fish were imaged in similar lateral position under both bright-field (white) and fluorescent light. The final representative images were generated by superimposing bright-field and fluorescence images using Adobe Photoshop. Significance was assessed by Student’s t test.

### EdU staining

5-ethynyl-2’-deoxyuridine (EdU, Molecular Probes, Life Technologies) was dissolved in DMSO to make a 10mM stock solution. This stock solution was further diluted 50 times using PBS to 200 μM, and 0.15 μl and 0.3 μl was injected into juvenile and older zebrafish, respectively. After 6 hours EdU treatment, tumor fish were euthanized with an overdose of MS-222 (Tricaine) and fixed in 4% paraformaldehyde (PFA) at 4°C overnight. Then, the fixed samples were soaked using 25% sucrose (in PBS) at 4°C overnight. Finally, the samples were embedded in OCT medium and the medium was allowed to solidify on dry ice. Tissue was sectioned using a Leica CM1510 S Cryostat and tissue sections were placed on plus gold microscope slide (Fisherbrand). The slides were post-fixed in 4% PFA for 15 mins and permeabilized in 0.5% Trition x-100 in BPS for 20 mins. Then, freshly made Click-iT® Plus reaction cocktail was added to each slide and allowed to incubate for 30 mins. A small amount of Vectashield mounting medium with DAPI was added to the slide and covered with a coverslip. The EdU Click-iT Plus EdU Alexa Fluor 647 Imaging kit (Molecular Probes, Life Technologies) was used for EdU staining. Tissue sections were subsequently analyzed using an Olympus FV3000 confocal microscope. Significance was assessed by Student’s t test. This assay was completed using at least 3 biological replicates (multiple primary tumors).

### qPCR Analysis

Tumors were extracted from euthanized fish and homogenized inside a 1.5 ml Eppendorf tube using a pestle connected to a Pellet Pestle™ Cordless Motor (DWK Life Sciences Kimble™ Kontes™). Tumor RNA was extracted using NEB’s Monarch RNA mini prep kit. cDNA was synthesized from the extracted RNA using Invitrogen’s First Strand Synthesis cDNA kit. Following cDNA synthesis, real-time qPCR was performed in 384-well plates using Applied Biosystem’s Sybr Green qPCR MasterMix (Comparative Cт (ΔΔCт) method). Results were compiled and analyzed using the QuantStudio 7 Flex system (Applied Biosciences). PCR primers are provided in Table 1. All assays were completed using technical replicates, with at least 3 tumors tested per experimental group.

**Table 1:**
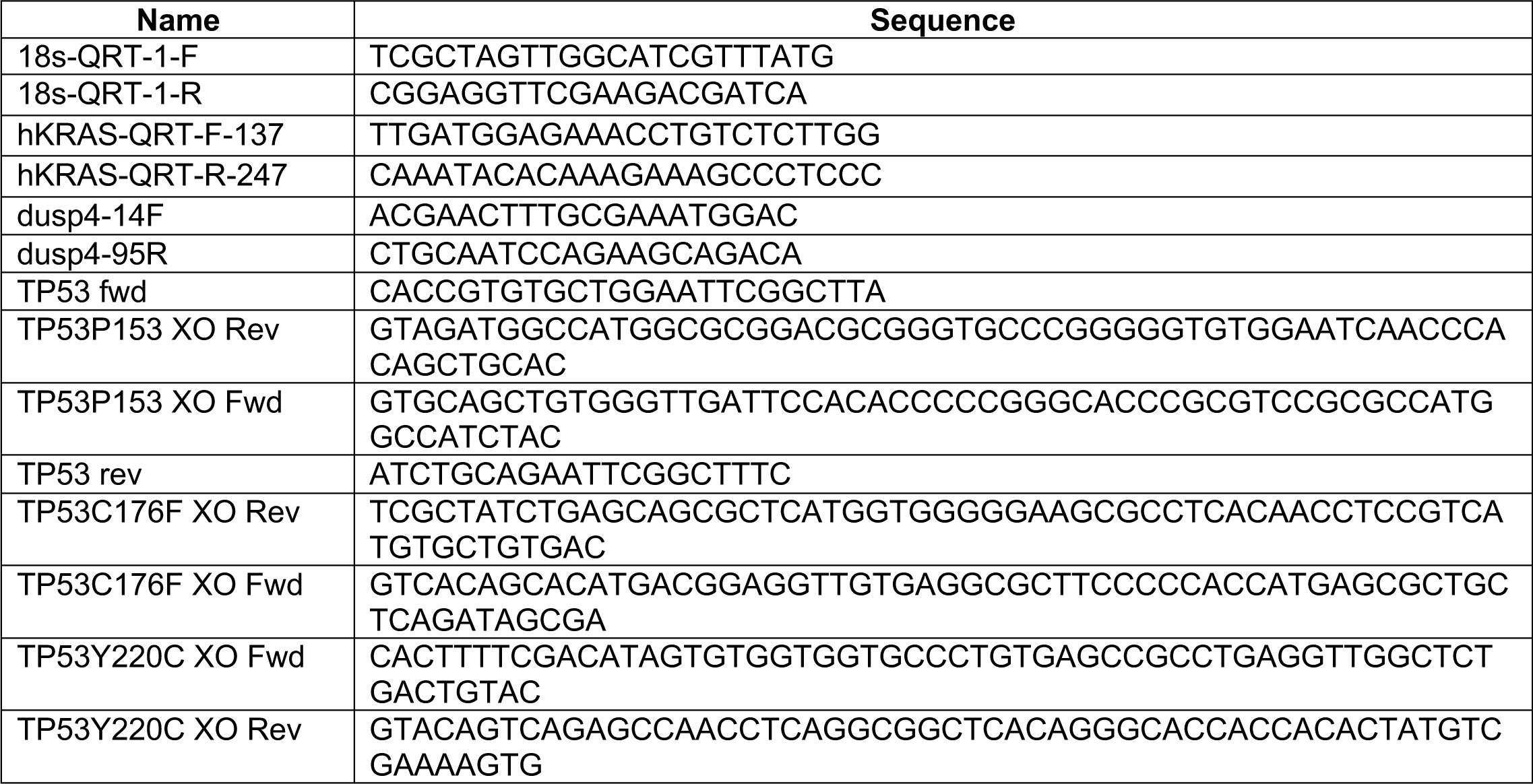
List of Primers used in this study.

**Table 2:**
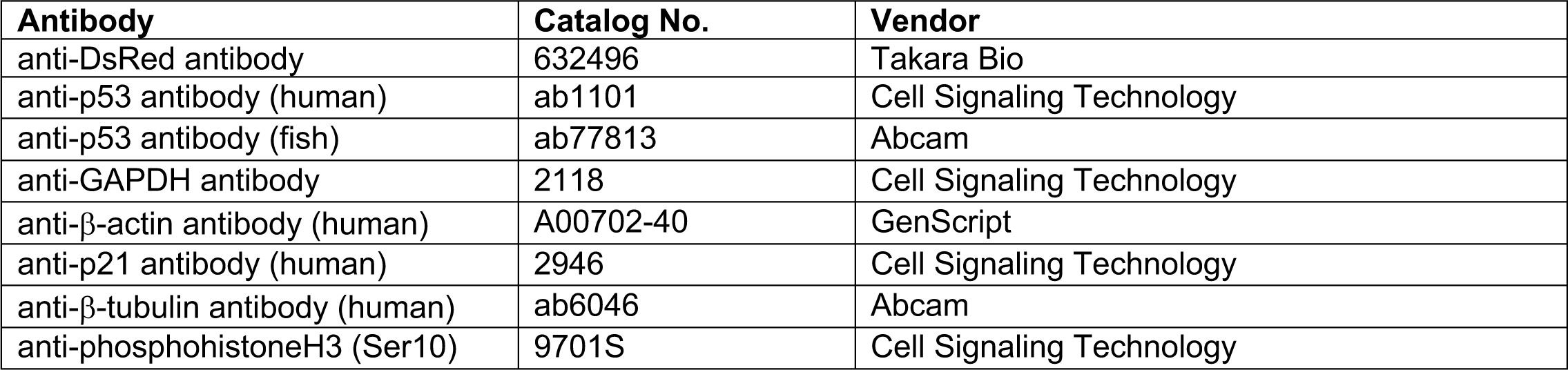
List of antibodies used in this study.

**Table 3:** List of reagents used in this study.

|  | Catalog No. | Vendor |
| --- | --- | --- |
| Click-iT™ EdU Alexa Fluor™ 647 Flow Cytometry Assay Kit | C10419 | ThermoFisher Scientific |
| Annexin V, Alexa Fluor™ 647 conjugate | A23204 | Invitrogen |
| IncuCyte® Caspase-3/7 Green Apoptosis Assay Reagent | 4440 | Sartorius |
| ZMC1 | HY-18634 | MedChemExpress |

### Phospho-histone H3 staining

Fish were fixed in 4% PFA at 4°C overnight. The fixed fish were subsequently soaked in 25% sucrose overnight and then embedded in OCT medium before being sectioned at 10 μm with a Leica CM1510 S cryostat. After being washed three times in PBST (0.1% Triton X-100 and 0.1% Tween 20 in PBS), the sections were incubated in blocking solution (2% horse serum, 10% FBS, 0.1% Triton X-100, 0.1% Tween 20, 10% DMSO in PBS) for 60 min. The sections were then incubated with rabbit anti-phospho-histone H3 (Ser10) primary antibody (1:500 dilution) at 4°C overnight. The following day, sections were washed three times in PBST and incubated with Alexa Fluor 647 conjugated anti-rabbit secondary antibody at room temperature for 2 hours. Vectashield mounting medium with DAPI was added to the slide and then a coverslip was placed over the sample. The slides were dried in the dark and sealed by nail polish. Sections were imaged using an Olympus FV3000 confocal microscope. Significance was assessed by Student’s T test. This assay was completed using at least 3 biological replicates (multiple primary tumors).

### Annexin V-FITC/PI staining

Fish were euthanized and tumor was isolated, following which the tumor was homogenized in 0.9x PBS + 5% FBS manually using a razor blade and made into single suspensions using 45 micron filters. Tumor cells were washed with 0.9x PBS + 5% FBS and resuspended in the binding buffer containing Annexin V-FITC and Propidium Iodide for 15 min in the dark at room temperature. Then the cells were detected by flow cytometry (FCM, FACS Canto™, BD, CA, USA). Significance was assessed by Student’s T test. This assay was completed using at least 3 biological replicates (multiple primary tumors).

### Histology and immunohistochemistry

Euthanized zebrafish were fixed in 4% PFA overnight at 4°C. Embedding, sectioning, and immunohistochemical analysis of zebrafish sections were performed as previously described (E. Y. Chen et al., 2014; Ignatius et al., 2012). H&E staining was performed at the Greehey CCRI histology core. Slides were imaged using a Motic EasyScan Pro slide scanner. Pathology review and staging were completed by board-certified pathologist (E.Y.C and A.R.G).

### Cloning *TP53* wild-type and mutants constructs

Wild-type *TP53* from both human (Addgene plasmid #69003) or zebrafish (3 days old embryos cDNA) were amplified by PCR and cloned into pENTR-D-TOPO vector, which was verified by DNA sequencing. *rag2-TP53* (human) or *rag2-tp53* (zebrafish) plasmids were generated by one-step Gateway reaction between a Gateway-compatible plasmid with the zebrafish *rag2* promoter flanked by *attR* sites and the respective pENTR-D-TOPO plasmid. *TP53^C176F^*, *TP53^P153Δ^*, or *TP53^Y220C^* fragments were constructed by amplifying human *TP53* as two separate fragments with the respective mutations, with the 3’ end of the first fragment possessing 60bp of homology with the 5’ end of the second fragment. These two fragments were purified from a 1% agarose gel using a Macherey-Nagel purification kit and spliced together using overlap extension PCR with Phusion^TM^ high fidelity DNA polymerase (Szymczak-Workman, Vignali, & Vignali, 2012). The entire spliced fragment was then blunt-ligated into a pENTR-D-TOPO vector. The *TP53* insert was sequenced after which it was cloned into a Gateway-compatible plasmid with the zebrafish *rag2* promoter flanked by *attR* sites using a one-step Gateway reaction using Gateway™ LR Clonase™ Enzyme mix. All other PCR amplification was carried out using Q5 high fidelity DNA polymerase.

### Western Blot Analyses

Western blot analysis on fish tumors was performed by first extracting tumor from fish and homogenizing tumor cells suspended in SDS lysis buffer inside a 1.5 ml Eppendorf tube using a pestle connected to a cordless motor. The total protein concentration for each lysate solution were normalized with a BCA assay kit (Thermo Fisher Scientific, Carlsbad, CA, USA). Then, 40 μg of total protein was run on a 10% SDS/PAGE gel. The protein transferred membrane were blocked using 5% fat free milk in TBST, followed by incubation in the appropriate antibody (anti-p53 antibody (1:1000 dilution in 5% BSA), anti-GAPDH (control, 1:1000 in 5% BSA), anti-p21 antibody (1:2000 dilution in 5% BSA).

### *In vitro* cell growth and colony formation assays

Gateway cloning was used to generate pCW57.1 with *TP53^WT^* or *TP53^P153Δ^* or *TP53^C176F^* or *TP53^Y220C^*, following which the *TP53* inserts were verified by DNA sequencing. These plasmids were transduced into a SaOS2 cell line via lentiviral infection using puromycin selection. The resultant puromycin-resistant stable line was treated with 2 μg/ml Dox for 2 days to induce *TP53* expression. Western blot analysis was carried out to check *TP53* expression in these cells. These cells were also seeded into 24-well plates at 20% confluency and stored at 37°C in the Incucyte ZOOM (Essen Bioscience). Cell confluency (%) was monitored every 4 hours over a period of 4 days. For colony formation assays, cells were seeded in 12 well plates (300 – 600 cells/well). After 24h, Dox was added to the wells at 2 μg/ml concentration. Incubation time for colony formation assays were around ∼2-3 weeks. When colonies were sufficiently large, media was gently removed from each plate by aspiration, and colonies were fixed with 50% methanol for 30 minutes at room temperature. Colonies were then stained with 3% (w/v) crystal violet in 25% methanol for 15 minutes at room temperature, and excess crystal violet was washed with dH2O with plates being allowed to dry. Colony formation was analyzed using ImageJ (Fiji). Significance was calculated by one-way ANOVA with Dunnett’s multiple comparisons tests. All assays were performed using technical triplicates, and at least 3 biological replicates were completed.

### Identifying *TP53* mutation status in Osteosarcoma PDX sample

DNA sample from osteosarcoma PDX was isolated using the Qiagen Puregene Core Kit A. PDX sample was first homogenized in lysis buffer, followed by heating at 65°C for 30 minutes. Thereafter, RNAse A was added to the sample and incubated for 30 minutes at 37°C. Protein Precipitation buffer was used to precipitate protein, and the sample was vortexed and then centrifuged at 15,000 RPM for 3 minutes. The supernatant was then removed, and isopropanol was added to precipitate genomic DNA, following which the sample was centrifuged at 15,000 RPM for 2 minutes at 4°C. The supernatant was then drained, and the DNA pellet was washed with 70% ethanol, followed by centrifugation at 15,000 RPM for 1 minute. The DNA pellet was then resuspended in DNA hydration solution, incubated at 65°C for 1 hour to dissolve DNA, and then incubated at 22°C for 1 hour. This precipitated DNA was used as a template in a PCR reaction using *TP53-*specific primers flanking the A159 and P153 residues. The PCR amplicon was ligated into a pCR™4-TOPO™ blunt cloning vector, and the ligation mix was transformed into DH5*α* chemically competent cells. Plasmid DNA was isolated from transformed colonies and sequenced using M13 primers.

### Luciferase assay

Stable SaOS2 cell lines harboring pCW57.1 plasmid (either empty vector or pCW57.1 with the appropriate *TP53* allele cloned into it) were first treated with 2 μg/ml Dox for 2 days to induce *TP53* expression, which were then transfected with a combination of PG13-luc assay plasmid and a control plasmid pRL-SV40P (50:1) using Lipofectamine 2000 (Invitrogen, USA) in accordance with the manufacturer’s instructions. After 48h of transfection, cell lysates were prepared and firefly and Renilla luciferase activities were measured by using the Dual-luciferase assay system according to the manufacturer’s instructions (Promega). All values are mean ± SD from at least three independent experiments. Significance was calculated by one-way ANOVA with Dunnett’s multiple comparisons tests. This assay was performed using technical triplicates, with 3 biological replicates completed.

### Caspase Glo 3/7 assay

SaOS2 cells were seeded at 20% confluency in 24-well plates and placed in the Incucyte, with half of the wells treated with 2 μg/ml Dox for two days to induce p53 expression. After reaching ∼40% confluency, media was supplemented with Caspase-Glo 3/7 reagent (1:1,000, Essen Bioscience) and Nuclight reagent (1:500, Essen Bioscience). Images of wells with and without Dox treatment were acquired at 24h post Caspase 3/7 in well-staining and processed using Adobe Photoshop and analyzed using ImageJ Cell Counter to determine percent caspase 3/7 events. Significance was determined using a two-way ANOVA with Sidak’s multiple comparisons test. This assay was performed in biological triplicates, using 3 technical replicates per assay.

### ZMC1 treatment

Tumor-bound fish were incubated in fish water containing ZMC1 at 70 nM concentration with another batch of tumor-bound fish incubated in 0.1% v/v DMSO (diluted in fish water) as control. Fish were imaged every week on an Olympus MVX10 stereomicroscope with an X-Cite series 120Q fluorescence illuminator and size of tumors was measured from the acquired images using FIJI software.

### Homology Modeling

The p53^P153Δ^ mutant was modeled using the SWISS-MODEL (Schwede, Kopp, Guex, & Peitsch, 2003) homology modeling server. The mutant p53 sequence and the p53^WT^ crystal DNA-binding domain crystal structure 2XWR (Natan et al., 2011) were used as input files. The homology models are built, scored and selected using statistical potentials of mean force scoring methods. Side-chain rotomers for non-conserved residues are selected through a local energy minimization function and the final models are subjected to global energy minimization using the CHARM22/CMAP forcefield (Waterhouse et al., 2018).

## Author contributions

JC and KB performed and/or interpreted or supervised aspects of the different experiments and helped write the manuscript; NH, LW, AL, AC, DL and AB helped with different experiments and analysis; AB and DSL provided structural analyses for the p53^P153Δ^ mutant. EYC and AG both soft tissue pathologists, helped with histology and immunohistochemistry analysis. PH, GT, AS, and AL discussed the clinical importance of the results. MI provided overall study direction, funding, and writing of the manuscript. All authors critically reviewed the report and approved the final version.

## Supplemental Figures

**Figure S1:**
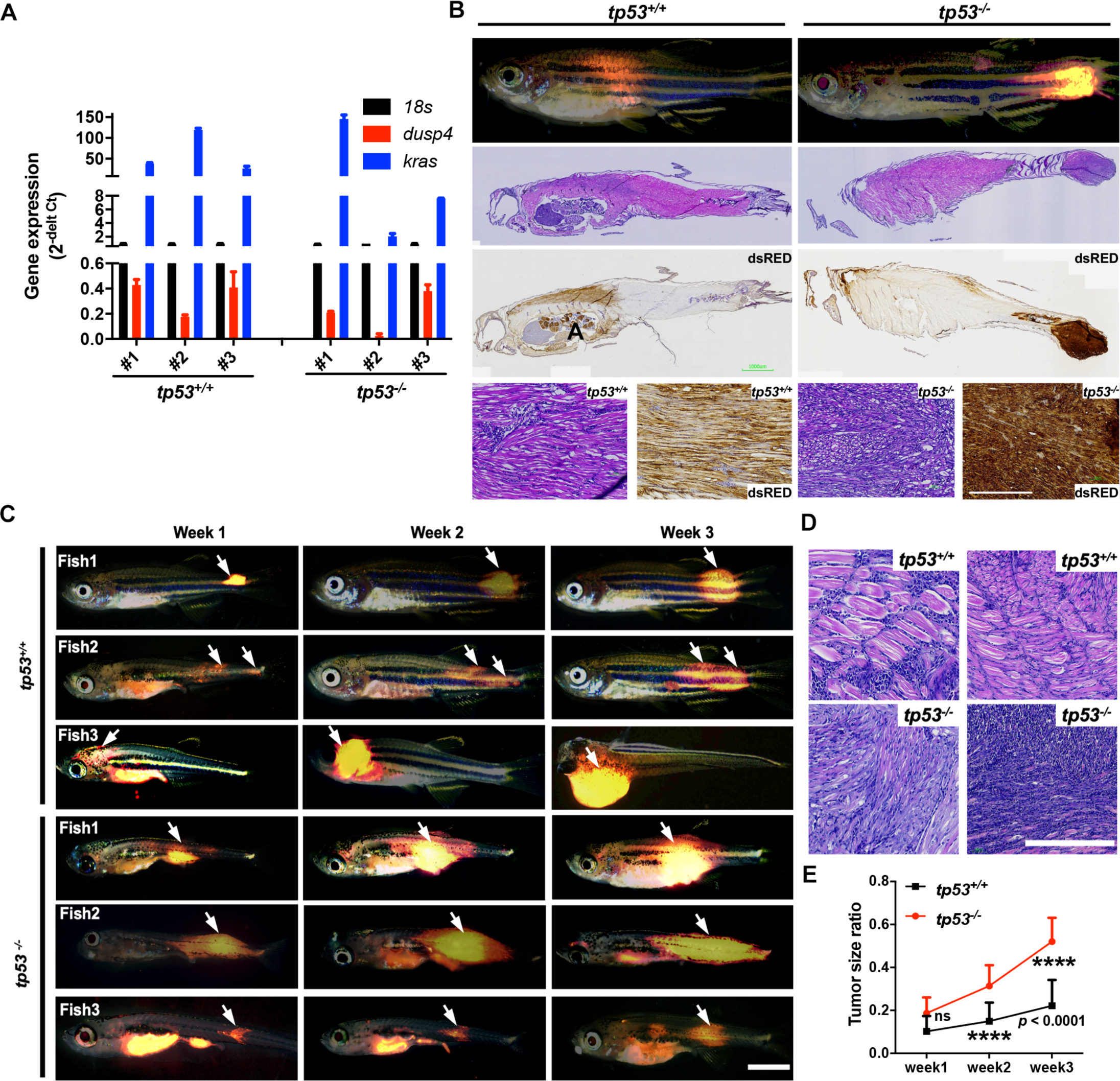
(A) mRNA expression of *kRAS^G12D^* and *dusp4* in compared to *18s* expression level. (B) Representative H&E and immunohistochemistry (anti-DsRed) staining in ERMS tumor fish. Arrows denote location of tumors. (C) Assessment of tumor growth and proliferation over 3 weeks in *tp53^-/-^* and *tp53^+/+^* fish (3 representative fish per group). Scale bar = 0.5 mm. (D) Representative H&E staining of *tp53^-/-^* and *tp53^+/+^* tumors. Scale bar = 100 µm. (E) Tumor size ratio per animal calculated over 3 weeks for *tp53^-/-^* and *tp53^+/+^* fish. ns = not significant. n = 13-16.

**Figure S3:**
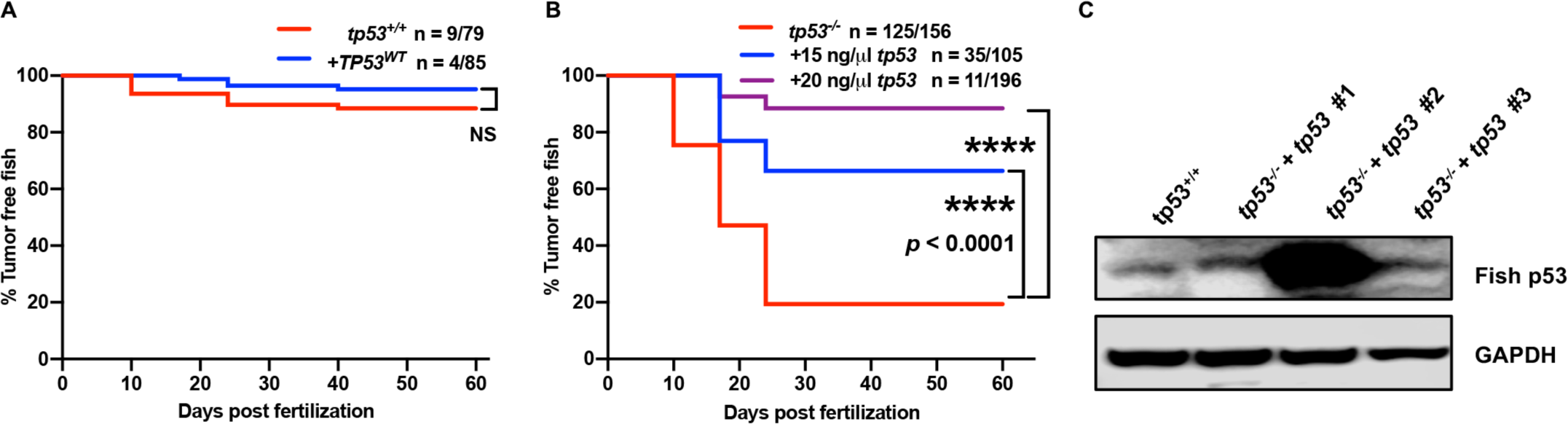
(A) Kaplan-Meier plot showing ERMS tumor initiation in *tp53^+/+^* fish (AB WT strain) with or without *p53^WT^*. ns = not significant (B) Kaplan-Meijer plot showing ERMS tumor initiation in *tp53^-/-^* fish (AB WT strain) with or without *tp53*. (C) Zebrafish p53 expression levels in three *tp53^-/-^* tumors expressed rag2-*tp53* along with rag2-*kRAS^G12D^*, using AB WT tumor driven by rag2-*kRAS^G12D^* as control.

**Figure S4:**
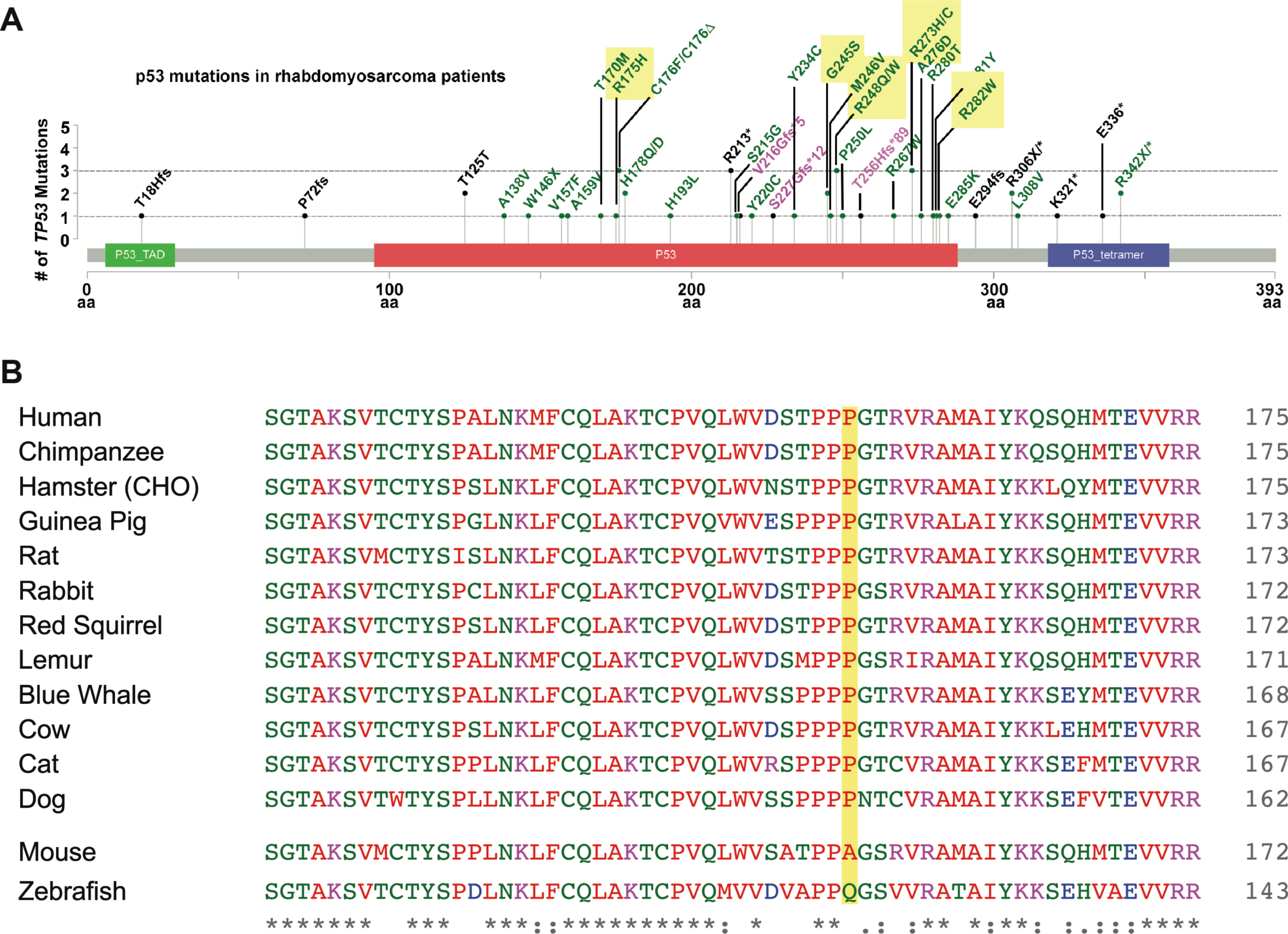
(A) Lollipop plot showing common *TP53* mutations in Rhabdomyosarcoma patients as well as the seven most common *TP53* hot spot mutations across all cancers. (B) The amino acid sequence alignment for *TP53*^P153^ with other species. The yellow highlight the amino acid of P153, and the top panel showed the conserved ones and bottom five species showed the non-conserved ones.

**Figure S5:**
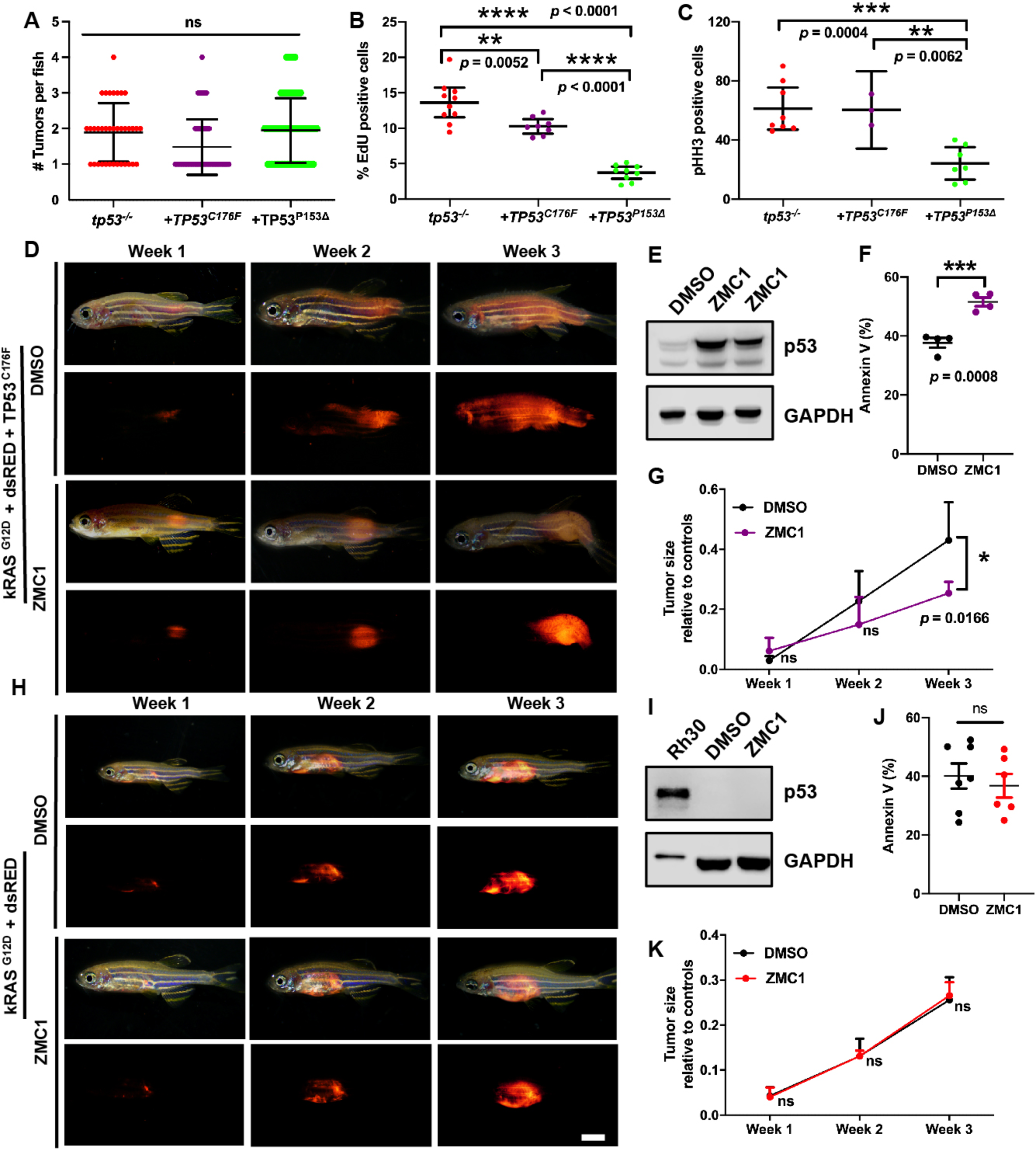
(A) Tumor numbers per zebrafish expressing no *TP53 (tp53^-/-^)*, or *TP53^C176F^*, or *TP53*^P153Δ^. n = 36 (*tp53^-/-^*), n = 58 (*TP53^C176F^*), n = 81 (*TP53*^P153Δ^). (B) Percentage of EdU positive cells in representative areas of fish tumors expressing either no *TP53 (tp53^-/-^)*, *TP53^C176F^*, or *TP53*^P153Δ^. n = 8-10. (C) Number of phospho-histone H3 positive cells in representative areas of fish tumors expressing either no *TP53 (tp53^-/-^)*, *TP53^C176F^*, or *TP53*^P153Δ^. n = 3-8. (D) Representative images of tumor burdened zebrafish with tumors expressing *kRAS^G12D^* + *TP53^C176F^* (DsRed+) treated with either DMSO or ZMC1 over a span of 3 weeks. (E) Western Blot showing p53 expression levels in fish tumors expressing *kRAS^G12D^* + *TP53^C176F^* and treated with either DMSO or ZMC1. (F) Percentage of Annexin V positive cells in tumors expressing *kRAS^G12D^* + *TP53^C176F^* that were treated with either DMSO or ZMC1. n = 4. (G) Ratio of tumor size to total body for in fish tumors expressing *kRAS^G12D^* + *TP53^C176F^*, treated with either DMSO or ZMC1. ns = not significant, n = 4. (H) Representative images of tumor burdened zebrafish expressing *kRAS^G12D^* (DsRed+) treated with either DMSO or ZMC1 over a span of 3 weeks. (I) Western Blot showing p53 expression levels in Rh30 cells (control) and fish tumors expressing *kRAS^G12D^*, treated with either DMSO or ZMC1. (J) Percentage of Annexin V staining of fish tumors expressing *kRAS^G12D^*, treated with either DMSO or ZMC1. ns = not significant. n = 6-7. (K) Ratio of tumor size to total body in tumors expressing *kRAS^G12^*, treated with either DMSO or ZMC1. ns = not significant, n = 4.

**Figure S6:**
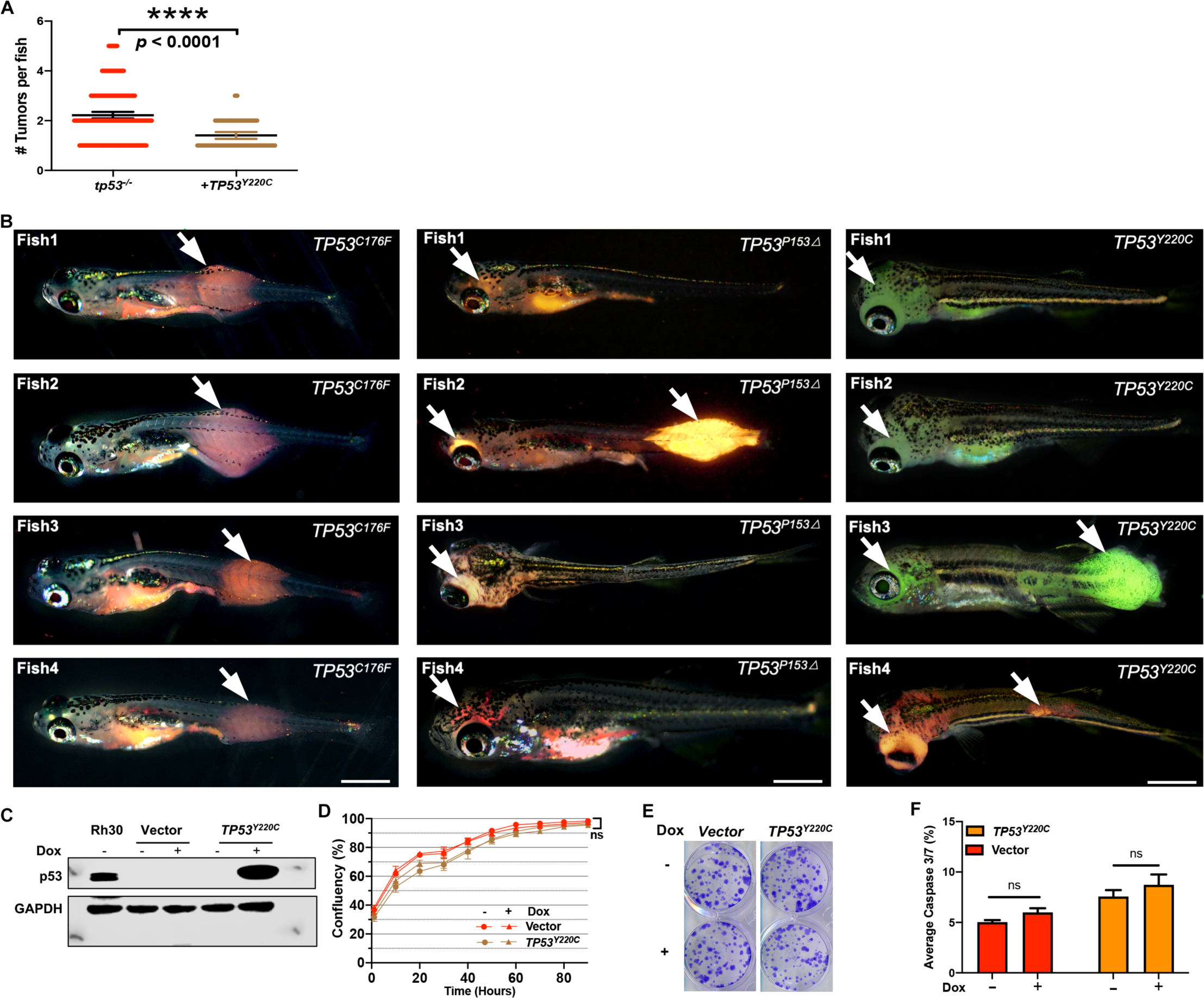
(A) Tumor numbers per zebrafish tumor expressing either no *TP53 (tp53^-/-^)*, or *TP53^Y220C^*. n = 89 (*tp53^-/-^*), n = 54 (*TP53^Y220C^*). (B) Representative images showing variation in tumor localization in fish expressing mutant *TP53* (either DsRed positive or GFP positive). (C) Protein expression of p53^Y220C^ expressed from a dox-inducible pCW57.1 vector in SaOS2 cells, using Rh30 as a positive control. (D) Growth curves (% confluence) of SaOS2 cells harboring dox-inducible pCW57.1 vector with or without *TP53^Y220C^* in the presence and absence of dox. ns = not significant (E) Colony formation assay for SaOS2 cells harboring dox-inducible pCW57.1 vector with or without *TP53^Y220C^* in the presence and absence of dox. (F) Quantification of caspase 3/7 assay to measure apoptosis for SaOS2 cells harboring dox-inducible pCW57.1 vector with or without *TP53^Y220C^* in the presence and absence of dox. ns = not significant

